# High-fidelity rare structural variant detection with HiFiRE3 reduced representation via restriction enzyme ends

**DOI:** 10.64898/2026.06.24.734375

**Authors:** Joseph A. Stewart, Jeanmarie Mishler, Samreen Ahmed, Bjoern Schwer, Thomas W. Glover, Thomas E Wilson

**Affiliations:** Department of Pathology, University of Michigan, Ann Arbor, MI, USA; Department of Human Genetics, University of Michigan, Ann Arbor, MI, USA; Department of Neurological Surgery, University of California, San Francisco, San Francisco, CA, USA; Department of Cellular & Molecular Pharmacology, University of California, San Francisco, San Francisco, CA, USA

## Abstract

High-fidelity detection of rare structural variants (SVs) remains challenging because library preparation and sequencing techniques generate artifactual junctions that obscure true single-molecule events. Here, we present HiFiRe3, an error-minimized sequencing framework that combines artifact-aware library design with error suppression and correction strategies to enable rare SV detection and frequency assessment across long and short-read sequencing platforms. We first systematically characterized major classes of SV artifacts, including chimeric PCR products, intermolecular ligation, sequencing platform-specific artifacts, and mapping errors. HiFiRe3 supports error correction of these artifact junctions by combining reduced representation restriction fragments with pre-ligation size selection to enable computational filtering via independent forced restriction enzyme end (FREE) and <1N size logics. In nanopore libraries, these approaches enabled targeted detection of single-molecule SVs at replication stress hotspots in cultured human cells exposed to genotoxicants and in long genes in untreated mouse brains, while markedly reducing singleton translocation artifacts. HiFiRe3 extends to PacBio sequencing for joint SV and SNV error correction and to short-read platforms for cost-efficient high-fidelity nonhomologous SV analysis. Together, HiFiRe3 is a flexible framework for accurately detecting rare genomic structural variation with broad applicability to targeted and genome-wide studies by selective application of its error correction approaches.

## Introduction

Structural variants (SVs) are a major source of genomic diversity and disease-associated change (1–3). They can alter gene dosage, disrupt coding sequences, reshape regulatory landscapes, and create novel rearrangements that are invisible to single nucleotide variant (SNV)-focused analyses. Detecting rare SVs accurately is especially important in settings such as cancer, normal tissue mosaicism, and *in vitro* mutagenesis, where true events of interest may exist at very low frequencies and can easily be masked by technical noise. This makes it essential not only to maximize sensitivity, but also to understand the specific artifacts introduced during library preparation, sequencing, and alignment so they can be suppressed or filtered.

Duplex sequencing was one of the earliest and most influential error-corrected sequencing strategies, developed to measure ultra-rare SNV frequencies by treating each strand of a DNA duplex as two independent measurements of the same sequence (4). In this method, adapters with random molecular tags are attached to both strands, the DNA is amplified and deeply sequenced, and reads are grouped by their tags to build single-strand consensus sequences before comparing the two complementary strands. Only variants confirmed on both strands are accepted as true SNVs. Duplex sequencing exploits the fact that real biological mutations appear in both DNA strands, whereas most PCR or sequencing errors arise on only one strand. Duplex sequencing is important for applications such as estimating *de novo* SNV rates in genetic toxicology (5,6), cancer genomics (7), and studies of DNA repair (8,9), where standard sequencing error rates would otherwise obscure the accurate measurement of rare mutation frequencies.

Many refinements of duplex sequencing have the same goal of extreme SNV accuracy, including CODEC, SMM-seq, ppmSeq, and others (10–12). Among them, NanoSeq and HiDEF-seq most directly influence work reported here (13,14). NanoSeq improved handling of damaged DNA to prevent errors arising during DNA end repair by cutting input DNA with blunt restriction enzymes (REs) and including ddBTPs during A-tailing (13). HiDEF-seq, for Hairpin Duplex Enhanced Fidelity Sequencing, combines independent readout of each DNA strand on the PacBio platform with library-preparation steps again designed to suppress artifacts arising during enzymatic handling (13). Smaller fragments than a typical PacBio library yield more passes of each strand in a closed covalent molecule for maximal accuracy. NanoSeq enabled ultra-accurate detection of somatic mutations at single-molecule resolution, with error rates reported below 5 errors per billion calls, making it possible to study mutation rates and signatures in diverse tissues (15). HiDEF-seq’s extreme fidelity allowed detection of both double-strand mutations and precursor single-strand DNA changes, helping reveal the initiating lesions behind mutagenesis (14).

While duplex technologies established robust SNV error-correction strategies, rare SV detection remains less well developed, which reflects the fact that the dominant error sources for SNVs and SVs are different (16). SNV calling is mainly affected by spontaneous or damage-induced base misincorporation during sequencing, whereas SV calling is more vulnerable to library preparation artifacts such as template switching and unintended ligation events that generate false junctions. In recent years, many methodological advances focused on improving SV calling rather than on library-based error correction. Long-read and assembly based callers such as VolcanoSV, cuteSV, SVIM, and Sniffles2 as well as short-read ensemble pipelines such as GATK-SV, Manta, and GRIDSS, are designed to reduce false positives by integrating split-read, discordant-pair, read-depth, and phasing evidence to resolve complex rearrangements based on reproducible detection over many reads (17–23). Such algorithms are distinct from error-minimization approaches that exploit knowledge of how a library was made to call single-molecule SVs analogous to duplex sequencing.

Some progress has been made in library-based error suppression and correction. SMM-SV-seq builds on the same single-molecule error-correction logic as SMM-seq based on rolling-circle amplification plus hairpin adapters with unique barcodes but adapts the workflow to detect somatic SVs in normal cells and tissues (11) [https://doi.org/10.1101/2024.08.08.607188]. We previously created svCapture, a targeted, probe-based enrichment approach that combines tagmentation-based libraries with over-sequencing of DNA inserts to establish strand family sizes in a manner analogous to duplex sequencing (16). As a ligation free approach, svCapture suppresses ligation artifacts while allowing error correction of chimeric PCR artifacts based on observations that PCR artifacts generally arise in later PCR cycles and thus have smaller strand families. svCapture enables highly specific detection of single-molecule SV junctions in target regions and has supported recent studies that established M-phase specific formation of SVs in synchronization experiments where only one copy of each unique SV-containing molecule is present (24). However, svCapture relies on costly application-specific probe sets targeted to small genomic regions and requires over-sequencing for error correction. A more generalized and less expensive approach would enable more studies.

Here, we report SV error correction approaches collectively called <u>Hi</u>gh <u>Fi</u>delity sequencing via <u>Re</u>striction enzyme-mediated <u>Re</u>duced <u>Re</u>presentation (HiFiRe3). HiFiRe3 is an adaptable combination of a library preparation protocol and a comprehensive computational tool suite that independently exploits RE-digested and size-selected DNA fragments for error-minimized sequencing focused on achieving high SV accuracy without over-sequencing. These reduced representation libraries can be prepared for both long and short-read sequencing platforms. The Oxford Nanopore-specific protocol allowed for flexible targeting and enrichment of larger regions of interest by adaptive sampling. We demonstrate the feasibility of measuring low frequency SVs in hotspot loci in RPE-1 cells, in recurrent DNA break cluster loci in mouse brains, and in response to bleomycin. Taking advantage of PacBio individual strand sequencing like HiDEF-seq, HiFiRe3 allows for error-minimized sequencing for both SVs and SNVs in the same reads. Together, HiFiRe3 is a high-fidelity sequencing approach for detecting rare genomic structural variation across platforms down to levels that support comparisons of single-molecule SV mutation rates analogous to SNV detection by duplex sequencing.

## Results

### SV error sources

In all high throughput sequencing data, downstream analysis can detect many SV junctions that did not exist in the source cells. These artifacts are generally recognizable as single-molecule supplementary alignments that mostly, but not exclusively, connect two random chromosomes. Our goal is to reduce these artifacts to a minimum so that the frequency signal from true rare variants – including those in a single sequenced molecule – rises above the background noise. HiFiRe3 achieves this by exploiting both error suppression, which minimizes artifact creation during library preparation, and error correction, which allows artifacts to be computationally filtered.

Common SV artifacts are mostly created by the enzymes used to add adapters to fragment ends but can also arise on the sequencer or during read alignment (Figure 1A). Artifacts arising during library preparation include polymerase template switching in PCR-mediated libraires and end-to-end ligation instead of adapter addition. Platform specific errors can cause two DNA inserts to appear as one read, such as a second strand entering a nanopore too quickly after a previous strand to be recognized as a new event. Such “follow-on” reads can occur between two strands from the same source molecule, appearing as a foldback inversion, or from different source molecules, appearing as any type of SV junction. Computational error sources include a reduced base quality, or just a low uniqueness, in portions of a read leading to incorrect supplementary alignments.

**Figure 1.**
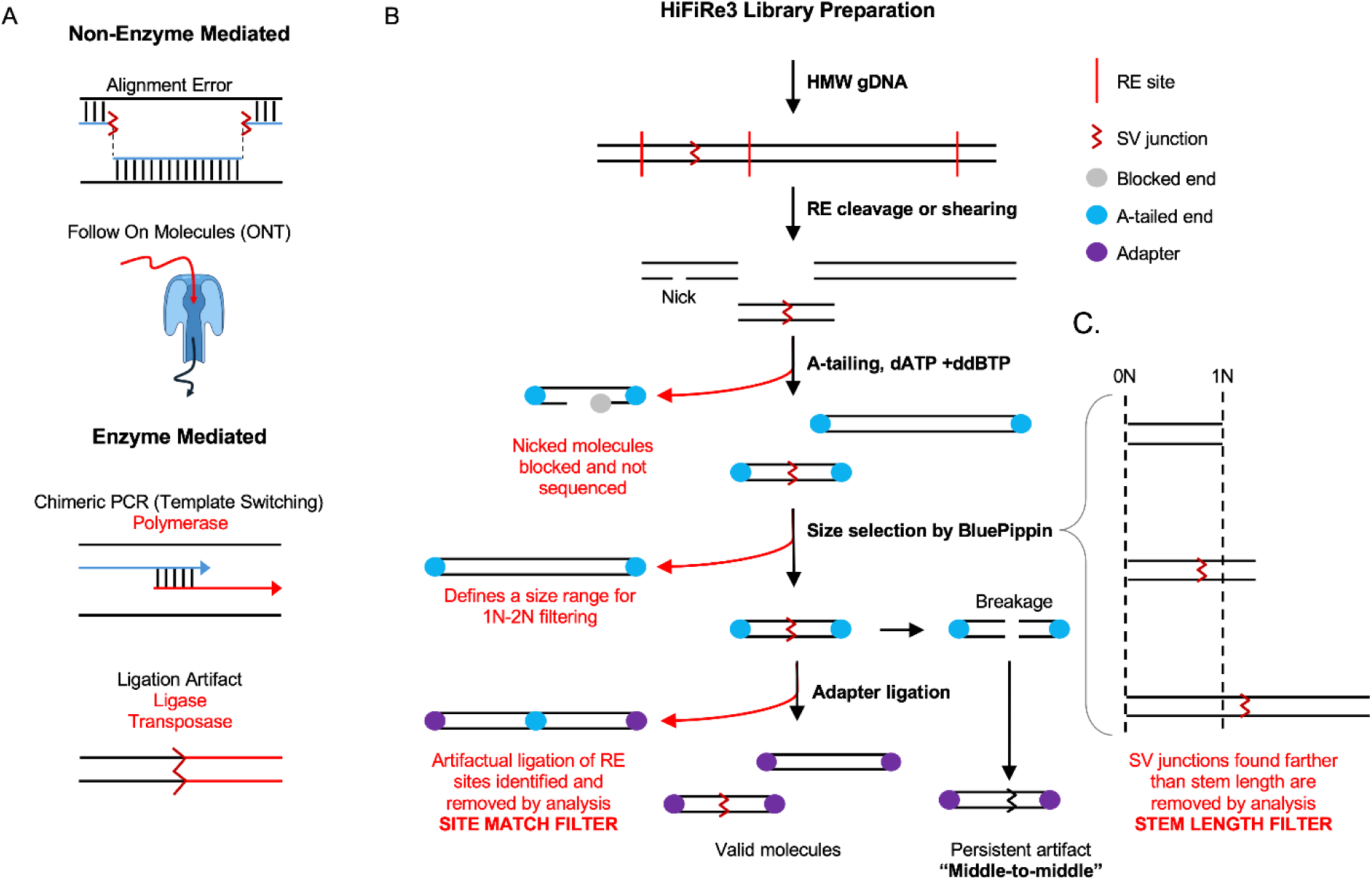
SV artifacts and HiFiRe3 library preparation scheme. **A.** Common SV artifacts can be categorized into non-enzyme and enzyme-mediated groups. **B.** The HiFiRe3 workflow starts with high molecular weight DNA and ends with a mixture of valid molecules and persistent artifacts. Different steps target specific artifact classes to make an error corrected library.

Understanding the contribution of these error sources in different library types allows the best error minimization strategies to be deployed, e.g., the HiFiRe3 pipeline handles ONT follow-on reads by rejecting junctions with stretches of low-quality bases. However, many principles below are generalizable. Enzymatic SV errors are best suppressed by not using the offending enzyme to prepare libraries. HiFiRe3 libraries are all PCR-free, which suppresses PCR chimera formation and promotes higher sequencing efficiency as compared to methods like svCapture that depend on over-sequencing. Unfortunately, all current sequencing platforms require an enzyme to add adapters. SV error correction demands knowing something about the nature of input DNA molecules to allow chimeras to be recognized, analogous to how the double-stranded nature of true SNVs enables duplex sequencing. Importantly, duplex logic cannot identify most SV artifacts in ligation-mediated libraries, which are inherently duplex events.

### HiFiRe3 error correction via forced restriction enzyme ends (FREE)

The first way HiFiRe3 enforces expectations about true source DNA molecules is by restricting where molecule endpoints are allowed to occur in a genome by digesting DNA with REs as in NanoSeq and HiDEF-seq (Figure 1B), which we refer to as “forced restriction enzyme ends” (FREE). Both *in silico* digestion of a genome and mapping of RE site polymorphisms based on observed read endpoints establishes *a priori* the genome positions at which DNA inserts are expected to end (Figure 2A). These RE sites are assigned to the base position in the reference genome immediately after the cleaved phosphodiester bond on the top strand (Figure 2B). SV junctions where either breakpoint mapped at or near a site position are rejected as chimeric by the SiteMatch filter against end-to-end chimeras (Figure 2D). Blunt-cutting REs help minimize SV and SNV artifacts by removing the need for multi-enzyme end repair in favor of A-tailing in the presence ddBTPs, which also blocks strand nicks.

**Figure 2.**
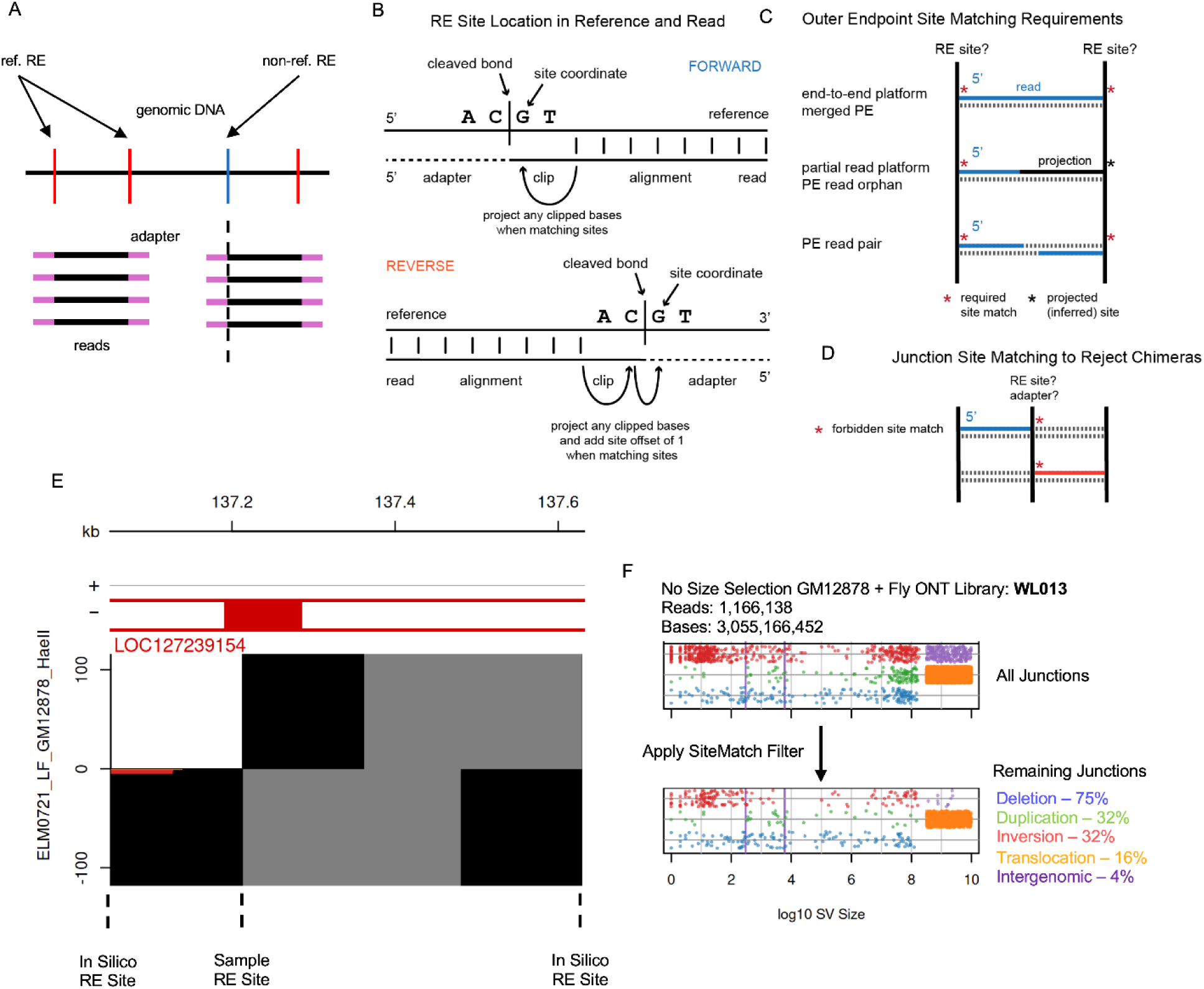
Restriction enzyme site determination and use in SV error correction. **A.** Genomic DNA is first digested *in silico* to determine RE sites (red) where adapters (pink) are expected to be added. However, if reads frequently begin or end in another location, a sample-specific RE site is added to the list used for SiteMatch filtering (blue). **B.** RE site positions are the clip-adjusted base position in the reference genome following the inferred cleaved phosphodiester bond, whether matching an alignment on the forward or reverse strand. **C.** In end-to-end platforms, reads (blue) always have two verified ends at RE sites (black vertical). In partial read platforms, only the 5’ end of the read must match an RE site. Reads that terminate early are projected (black horizontal) to the next RE stie when tabulating insert sizes. **D.** If a junction breakpoint maps to an identified RE site, the SV is filtered and rejected. **E.** Browser view showing two sites where reads (black horizontal) begin and end in an Aviti 2x150 library, showing the nature of reduced representation libraries and the contribution of a sample-specific RE site. **F.** Junction summary plots showing singleton SV size on the x-axis and type by color (one dot per junction, deletion = blue, duplication = green, inversion = red, intraspecies translocation = orange, interspecies translocation = purple) from an ONT library containing human and fly DNA. Applying the SiteMatch filter alone (bottom graph) rejects a large percentage of reads.

Important features are first that both true ends of sequenced DNA inserts are not always identified by a read. The 5’ end of a read at the adapter always matches one true end of the insert, but the 3’ end in a partial read platform or a paired-end read orphan may not. In these cases, 3’ read ends are projected to the next known RE site to infer the source RE fragment (Figure 2C). These projected molecules are used in combination with molecules with two verified ends to tabulate fragment size distributions for a library. Secondly, molecules broken on both strands at positions internal to RE fragment ends might ligate to each other to create SV “middle-to-middle” artifact junctions (Figure 1B). Because these artifacts escape the SiteMatch filter, it is important to use high quality DNA with minimal breakage. We achieve this by digesting high molecular weight (HMW) DNA with REs while it is still wrapped around the two large glass beads used in the NEB Monarch HMW DNA extraction kit, where RE digestion proved sufficient to release DNA for library preparation without the need for complex DNA hydration protocols.

### HiFiRe3 size selection and <1N error correction logic

A distinct property that can be known about input DNA molecules *a priori*, i.e., before adapter addition, is their size. Importantly, size selection for the purpose of SV error correction must be performed before adapter ligation, in contrast to most protocols where size selection after adapter addition helps optimize platform-specific sequencing yield. In HiFiRe3, DNA molecules are size selected using a device such as a BluePippin between two fragment lengths we refer to as 1N and 2N, e.g., between 8 kb and 16 kb as used in many nanopore libraries below (Figure 3A). The smallest true DNA fragments in a library (1N) dictate the smallest possible size of a chimeric ligation artifact (2N). While it works to reject SV junctions in inserts with an inferred physical size >2N, a more generalized way to exploit pre-ligation sizing demands that any passing SV junction be within 1N bp from a validated DNA endpoint, a value we call the “stem length”. Such a junction could not have arisen by joining of two molecules whose smallest input size was 1N. Although molecules larger than 2N bp can contain valid junctions that pass the StemLength filter, they waste bases in central read regions that cannot contribute to SV junction detection, making 1N to 2N bp size selection most efficient. Size selection on the BluePippin also supports immediate transfer of A-tailed DNA molecules directly from the device to adapter ligation reactions, which minimizes the post-ligation fragmentation that leads to middle-to-middle chimeras.

**Figure 3.**
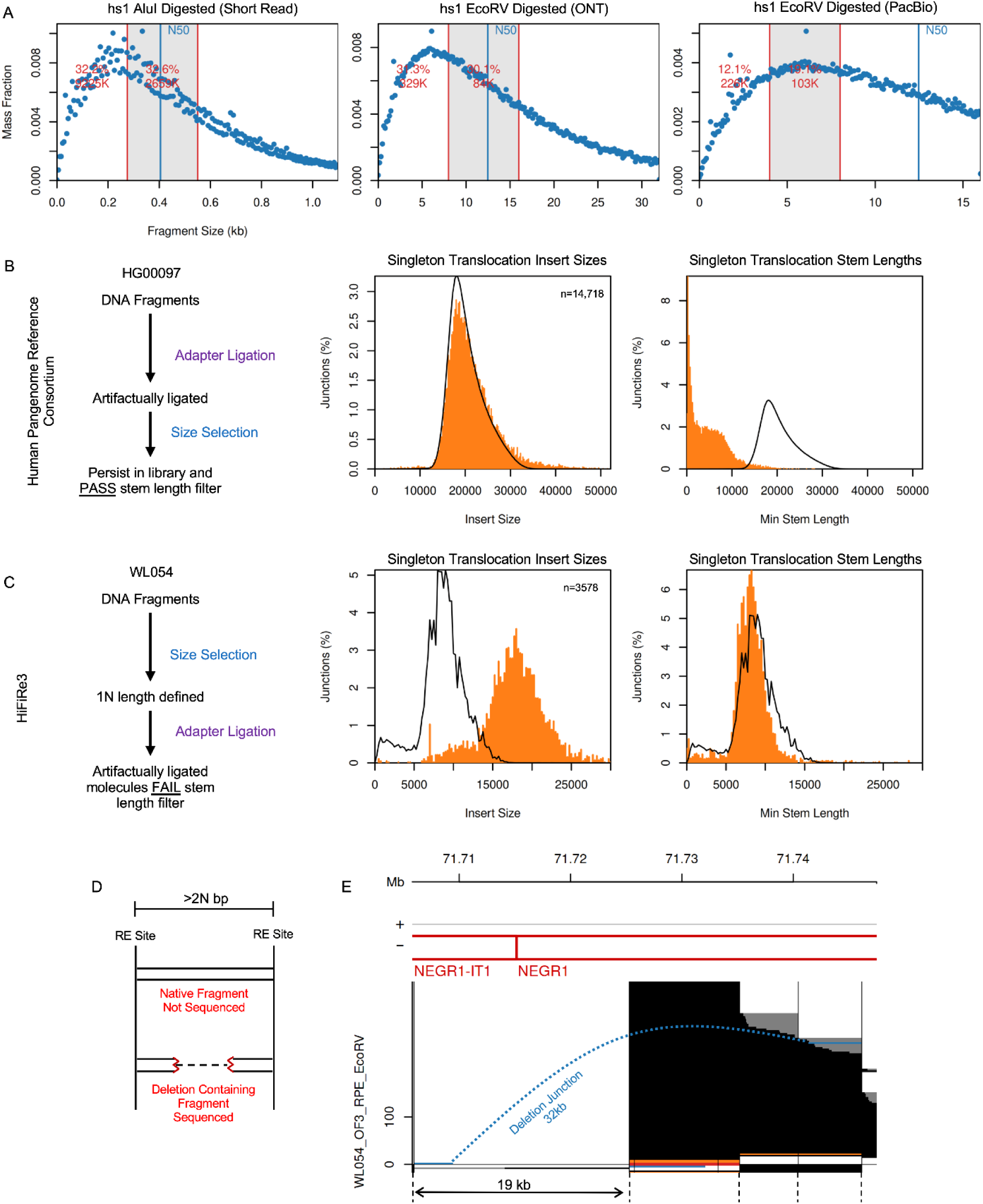
Size selection and stem length filtering for SV error correction. **A.** In silico RE fragment sizes of the hs1 genome by specified enzyme. The gray area between red vertical bars depicts the experimentally selected mass fraction. **B.** Schema (left) of a typical PacBio library prep from the Human Pangenome Reference Consortium, showing the frequency distribution of insert sizes (middle) and junction stem lengths (right) for non-SV molecules (black line) and singleton translocations (orange) as presumed artifacts. **C.** Like B, for HiFiRe3. Unlike B, artifact molecules are shifted to larger fragments >2N bp with stem lengths >1N bp. **D. and E.** Schema and browser view showing that SV-containing fragments, like non-SV fragments, will obey the size-selection limits, but need not arise from the same RE fragments, e.g., a deletion fragment (blue) breakpoint might arise in a fragment too large to otherwise sequence. RE sites are vertical black lines. Non-SV reads are black with projections in gray. Translocations are orange and inversions are red.

To illustrate the importance of performing size selection before adapter addition we compared HiFiRe3 fragment size distributions stratified by molecule type to a typical PacBio library created by the Human Pangenome Reference Consortium (HPRC) (25). The abundant SV artifacts in the HPRC library were necessarily created by the joining of two shorter molecules such that they made it through size selection after they were created. Artifact inserts thus, unlike HiFiRe3, have the same size distribution as true molecules and short stem lengths, which precludes size-based SV error correction (Figure 3B,C).

Importantly, HiFiRe3 can detect SV junctions anywhere in the genome with no bias for being in reference RE fragments within the selected size range (Figure 3D). For example, a junction breakpoint in a 20 kb RE reference fragment can be sequenced in a library selected for fragments 8 to 16 kb if the actual sequenced SV molecule was in the selected range. One deletion breakpoint in the *NEGR1* gene in Figure 3E is within a high coverage reference fragment that falls within the selected size range while the second is in a larger fragment that was not otherwise sequenced. The junction-containing molecule was sequenced because it fell in the selected size range.

### Intra-species vs. inter-species translocations for monitoring library quality

SV artifacts are expected to mostly join two random positions in different chromosomes, a fact we have exploited when assessing chimeric artifact rates (16). To be certain that singleton translocations are a valid means of tracking artifact rates and processes, we spiked *Drosophila* cells into human cells prior to HiFiRe3 DNA and library preparation and mapped reads to a composite reference of both species genomes. While intra-species junctions might sometimes be true source DNA molecules, SVs between two species must be artifacts. Figure S1 shows a non-size-selected, EcoRV-digested, ligation ONT HiFiRe3 library made from mixed GM12878 and *Drosophila* S2 cells at approximately 7% Drosophila DNA. It was adaptively sampled for 11 large genes making up a little over 1% of the human genome. HiFiRe3 reported 256 inter-species human-fly singleton translocation junctions and 4,827 singleton intra-species singleton translocations out of 1.16M total reads and 3.06B sequenced bp. We inferred the mechanisms that formed these junctions in part by calculating their “alignment offsets”, i.e., the number of (micro)homologous or *de novo* inserted bases, where negative alignment offsets are microhomologies and positive numbers are insertions. Both the intra-species and inter-species translocations showed peaks at alignment offsets of -6, -1, and a broader peak around 40 bp with similar patterns. This indicates that most intra-species and inter-species translocations are formed by the same mechanisms such that the more abundant intra-species events can be used as a reliable metric of library quality.

### Inferring SV error processes from observed junctions across platforms

Optimizing HiFiRe3 approaches required comprehensive assessment of the SV error classes that apply to different library types and sequencing platforms. The pipeline tracks all junctions revealed by supplementary alignments and groups them into distinct observed junctions in a manner that accounts for differences in breakpoint node assignment due to base errors in sequencing. Each final junction carries observation counts, alignment parameters such as mapping quality (MAPQ), and error correction metrics enabled by HiFiRe3 library methods.

In Figure 4A, we plot final junction calls for a nanopore ligation library where input DNA from aphidicolin (APH)-treated RPE-1 cells was fragmented using EcoRV, size-selected to 8 to 16kb, and subjected to adaptive sampling in large, expressed genes. All unfiltered junctions are plotted by their event size (x-axis), observation count, and junction type (y-axis). Most junctions are singleton translocations and presumed artifacts, in addition to a collection of small inversions and other deletions, duplications and inversions at varying sizes. The distribution of alignment offsets in these subsets showed a predominant insertion peak at around 35 bp (Figures 4C to E), the typical number of nanopore adapter bases sequenced in follow-on junctions, including foldback inversions. We also plotted the insert size distributions of these artifact groups compared to non-SV reads where artifact junctions had much larger insert sizes (Figures 4C to E). In contrast, less frequent small deletions mostly had larger insertions close to their deletion size that fail the TraversalDelta metric where stretches of low-quality bases give the false appearance of deletions with insertions between the breakpoints (Figure 4F and G). Since these molecules are not chimeric, their insert sizes are like those of non-SV reads. Because ONT follow-ons consistently carry a low-quality insertion, the LowQualIns filter in combination with SiteMatch and StemLength dominate the junction failure reasons (Figures 4C to F). Enforcing all filters leaves junctions with the highest likelihood of being a genuine SV (Figure 4B), which have a distinct microhomology profile compared to artifacts characterized by short microhomologies and small insertions typical of APH-induced end joining (Figure 4H).

**Figure 4.**
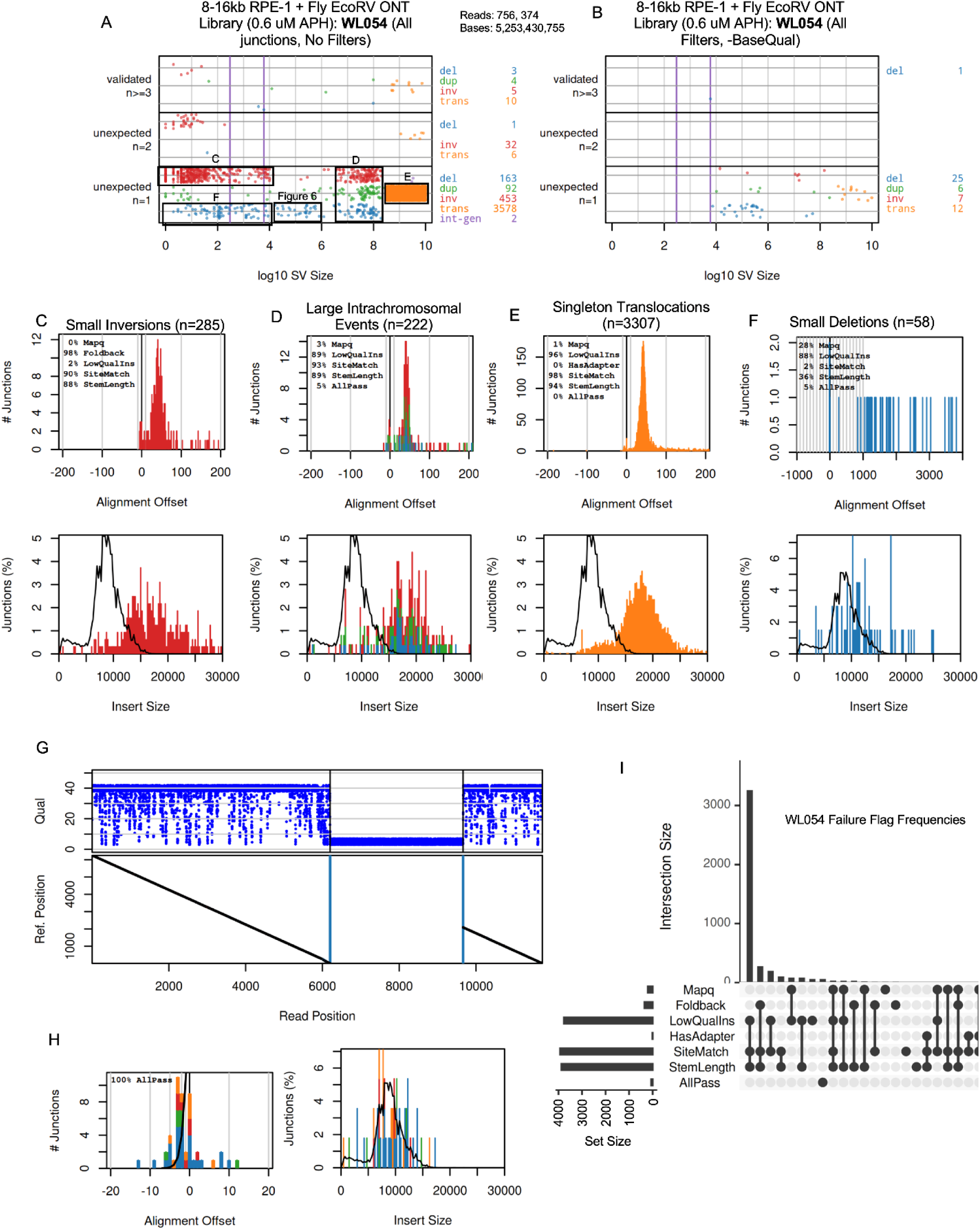
ONT library artifact summaries. **A.** Junction summary plot without any active filters showing SV size on the x-axis and type by color (one dot per junction, deletion = blue, duplication = green, inversion = red, intraspecies translocation = orange, interspecies translocation = purple) from an ONT library containing human and fly DNA. The y-axis depicts SVs called by a single read (bottom), two reads (middle), or three or more reads (top). Black boxes depict junction subsets used in alignment offset panel C to F. **B.** Like A, but with all filters active except BaseQual. **C. to E.** Histograms showing the number of junctions with insertions (positive alignment offset) or microhomology (negative alignment offset) (top) and insert sizes (bottom) for small inversions, large intrachromosomal events, and singleton translocations. Insets show junction failure percentages by filter type. **F. and G.** Alignment offsets for small deletions, and one example read that failed LowQualIns. Base quality (blue points) drops severely in the center of the read called as a deletion (vertical blue lines). Diagonal lines below show reference movement along the read. **H.** Alignment offset plot (left) and insert size plot (right) of remaining junctions when all filters (except MapQ for nanopore) are applied. **I.** Upset plot showing the relationship between different filtering flags for all library junctions.

In studies below, we also created non-size selected ONT libraries. Like the size-selected nanopore libraries, these libraries are prone to large insertions at their junctions but no longer have a clear separation in non-SV and artifact insert sizes (Figure 5A and B). Thus, most junctions in an ONT library are artifactual but HiFiRe3 can identify them independently of the observation count even without the benefit of size selection.

**Figure 5.**
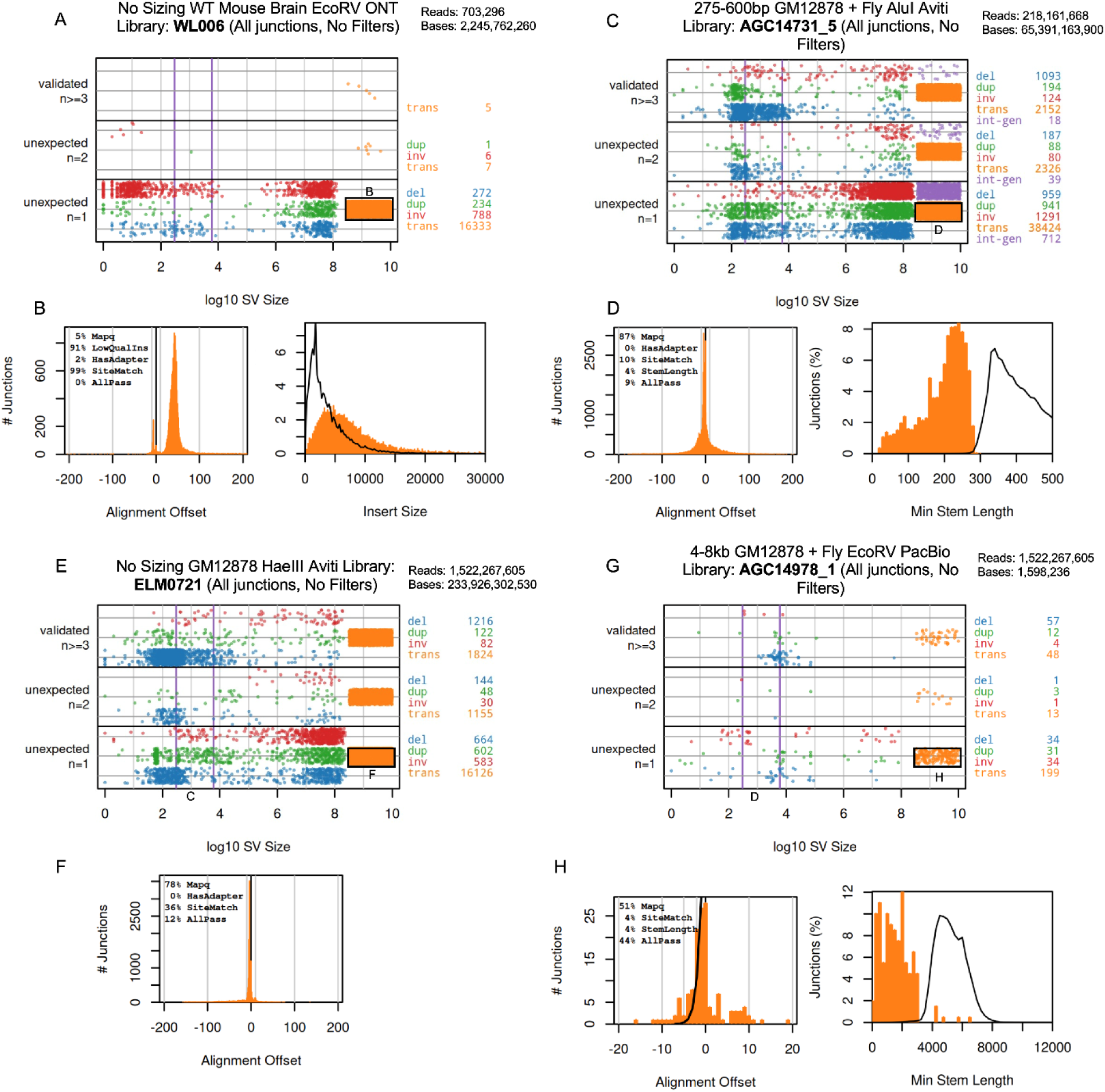
Comparative ONT, Aviti, and PacBio library artifact summaries. **A.** Junction summary plot like Figure 4A for a non-size-selected ONT library with no active filters. The black box denotes the read subset used in panel B. **B.** Alignment offset plot (left) and insert size plot (right) of singleton translocations from A with no filtering. **C.** Junction summary plot for a 1x300 Aviti library containing human and fly DNA with no active filters. The black box denotes the read subset used in panel D. **D.** Alignment offset plot (left) and minimum stem length (right) of singleton translocations from C with no filtering. **E.** Junction summary plot for a 2x150 Aviti library with no active filters. The black box denotes the read subset used in panel F. **F.** Alignment offset plot of singleton translocations from E with no filtering. **G.** Junction summary plot for a PacBio library containing human and fly DNA with no active filters. The black box denotes the read subset used in panel H. **H.** Alignment offset plot (left) and minimum stem length (right) of singleton translocations from G with no filtering.

While long-read sequencing has advantages for SV detection, short-read platforms are cost efficient and able to detect nonhomologous SVs in non-repetitive DNA. We explored short-read HiFiRe3 on the Element Aviti platform that supports higher base accuracy, avoids index hopping by using rolling-circle amplification, and supports 1x300bp single reads, all of which benefit SV analysis. Figure 5C shows a junction summary plot for an Aviti library made with GM12878 DNA cut with AluI, size-selected for fragments 275-600 bp and sequenced to 20X coverage as part of the bleomycin experiment below. Singleton translocations were again the most frequent SV class. Because Aviti follow-on molecules are not possible, these junctions are characterized by blunt ends (Figure 5D). Stem lengths for singleton translocations skewed to the right as they represent the leftmost tail of the artifact molecule size distribution. MapQ was by far the most frequently failed filter emphasizing that low confidence alignments must be properly handled in short-read libraries. We therefore replotted the data applying only the MapQ filter, such that both alignments flanking the remaining junctions were declared with high confidence (Figure S2). Those junctions mostly passed the rest of the filters, with most remaining failures coming from SiteMatch alone. Non-size-selected 2x150 paired-end libraries showed similar artifact trends (Figure 5E and F). Due to these libraries’ high, untargeted genome-wide coverage, validated junctions with three or more supporting reads were easily observed, many of which correspond to apparent deletions of Alu and LINE sequences relative to reference (vertical purple lines in all junction summary plots). Even after applying all filters, many potential singleton SVs remained, that showed a more restricted alignment offset profile (Figures S2B and G).

Finally, Figure 5G shows a junction summary plot for a PacBio library made with GM12878 DNA cut with EcoRV and size selected for fragments 4-8 kb, also part of the bleomycin experiment below. Much like Aviti libraries, most of the PacBio singleton junctions were caught by the MapQ filter. However, the stem lengths of these artifacts appear to skew smaller rather than larger (Figure 5H). This could be due to the low coverage in this library with 1.6B sequenced bases. An in-depth analysis of all artifact classes for each library type presented in this study can be found in Figures S2-S7.

### Inter-dependence of SV error correction filters

We created upset plots for nanopore (Figure 4I) and other HiFiRe3 libraries (Figure S2-S7) to explore the degree to which the various quality and error correction filters provide unique vs. redundant information. Each platform had a distinctive pattern of contributions from each filter. For ONT libraries, most failed junctions were caught by each of the SiteMatch, StemLength, and LowQualIns or Foldback filters (Figure 4I), again revealing that follow-on read fusions are the dominant error class. Non-follow-on/foldback events were most frequently caught by both SiteMatch and StemLength. These patterns reveal a significant non-independence of the different filters consistent with the fact that many chimeric molecules are expected to share multiple filterable properties. In contrast, StemLength rarely failed for Aviti libraries independently of MapQ (Figure 5F), consistent with the fact that the sequenced DNA inserts (275-600 bp) were mostly longer than the read length (300 bp) making it impossible to reach most end-to-end artifact junctions. Only SiteMatch was a significant independent contributor and most junctions passed all filters. Again, like Aviti, PacBio libraries are also dominated by MapQ alone as a failure reason followed by combinations of SiteMatch, StemLength, and Foldback filters. Together, these data show that HiFiRe3 is effective in both describing and correcting artifacts across a variety of sequencing technologies but that its error correction methods are often interdependent and need not all be applied (see Discussion).

### Identification of aphidicolin-induced SV hotspots in RPE-1 cells

To explore the ability of HiFiRe3 error correction to detect low frequency junctions in paradigms known to induce *de novo* SVs, we next examined common fragile sites (CFSs). CFSs are loci on metaphase chromosomes prone to forming gaps and breaks when subjected to replication stress (26). CFSs correspond to large, transcriptionally active genes, and so are cell type specific with respect to which genes are fragile. They are also known to be hotspots for copy number losses via intrachromosomal deletions with a median size of ∼200kb (27). We used HiFiRe3 with size selection and adaptive sampling for twenty-one transcriptionally active large genes in RPE-1 cells as candidate deletion hotspots, approximately 1% of the genome. We treated RPE-1 cells with either 0.6 μM or 0.4 μM APH, a B-family polymerase inhibitor known to induce replication stress, for 48 hours before harvesting cells and creating HiFiRe3 ONT libraries with 8-16kb fragments (Figure 6A).

**Figure 6.**
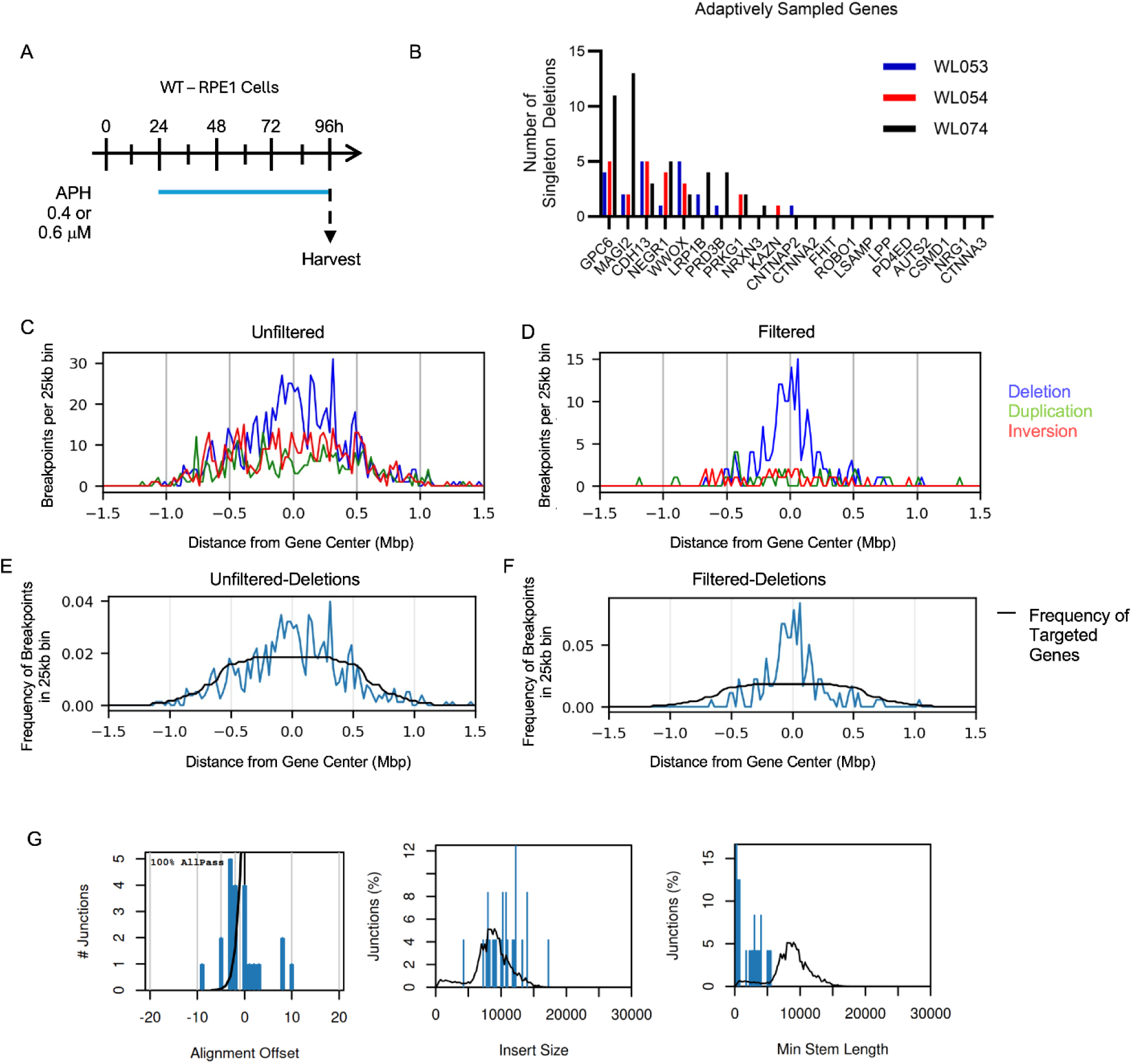
Identifying SV hotspots at large genes in RPE-1 cells. **A.** RPE-1 cells were grown for 72 hours in the presence of 0.4 or 0.6 μM APH before harvesting for library preparation. **B.** Number of called deletions per gene targeted by nanopore adaptive sampling over three experimental replicates. WL053 and WL054 were treated with 0.6 μM APH while WL074 was treated with 0.4 μM APH. **C. and D.** Traces depicting the number of SV breakpoints in 25kb bins by SV type as a function of distance to the center of their respective gene, without (C) and with (D) error correction filtering. **E. and F.** Frequency distribution of deletion breakpoints from C and D, without (E) and with (F) error correction filtering. The black line is the frequency distribution of the number of genes targeted by adaptive sampling for the same bins. **G.** Alignment offset, insert size, and minimum stem length distributions of filtered singleton deletions from 10kb to 1 Mb as boxed on the junction summary plot in Figure 4A.

With 8-16kb fragments we achieved 16-fold on-target enrichment, three times higher than earlier non-size selected libraries that had an average fragment length of about 5kb. With a combined 461X on-target coverage over three replicate libraries, we identified 88 singleton deletions between 10kb and 1Mb across 11 genes with the three most frequent genes – CDH13, GPC6, and MAGI2 – responsible for 57% of junctions (Figure 6B). The other 10 genes had no called SVs in any replicate. Before error correction filtering, breakpoints for all three intrachromosomal SV types were found randomly throughout the gene bodies at a density that mostly matched the coverage of targeted genes (Figure 6C, E, S8). Some weak enrichment for central deletions could be appreciated, but after error correction the deletions formed a strong, clear peak at the center of their respective genes (Figure 6D, F). They also had short microhomologies and small insertions typical of the POLQ-mediated end-joining process active at CFSs (24) and insert sizes that match non-SV reads and stem lengths <1N bp (Figure 6G). The fact that *de* novo deletions with expected structures were strongly biased to being in the centers of only a subset of the targeted genes matches the well-described biology at CFSs and validates the specificity of called junctions (16,24). Interestingly, duplication junctions, although less numerous, followed the opposite trend with enrichment on gene flanks following error correction (Figure S8), again consistent with prior observations (27). Thus, HiFiRe3 quickly and efficiently established CFS-associated SV hotspots in a cell line for which no prior data existed.

### Sensitive detection of SVs in large genes in mouse brain tissue

To test the ability of HiFiRe3 to detect rare *de novo* SV junctions *in vivo*, we focused on large genes in mouse brain tissue. Previous studies reported that large, neuron-specific genes are hotspots for DSBs referred to as recurrent DNA break clusters (RDCs) (28,29). RDCs were identified in neural stem/progenitor cells (NSPCs) from mice that lacked Xrcc4 and p53 using high-throughput, genome wide translocation sequencing (HTGTS). Interestingly, half of the identified RDCs overlapped with DSBs seen in WT NSPCs treated with APH, highlighting their similarity to CFSs. Here, we have begun to test the hypothesis that RDCs will also correspond to regions of somatic SV accumulation in mouse brain tissue.

We collected mouse brains from three different sources in two genotypes, WT and p53^-/-^ (Figure 7A). Brain tissue was dissected into hippocampus, cortex, or whole brain. HiFiRe3 libraries were prepared using the ONT ligation kit without size selection, with cleavage by EcoRV digesting the mouse genome on average into 5kb fragments. Nanopore sequencing adaptively sampled ten RDC genes representing approximately 1% of the mouse genome. We sequenced 17 mouse brain libraries across both genotypes and five mouse sources with 670X on-target coverage for WT and 965X coverage for p53^-/-^. The average on-target enrichment was approximately 5-fold. We detected SVs in all target genes, for example a deletion in *Wwox* in Figure 7B. The libraries had an average frequency of 6.7x10^-9^ singleton translocations per on-target sequenced base as an estimate of the SV error rate, with most libraries behaving similarly (Figure 7C). We identified a total of 231 singleton SVs in WT and 181 in the p53^-/-^ brain tissue, with translocations making up 74% of the SVs. To assess which of these SVs showed a random artifact vs. non-random junction pattern, we used one of the two breakpoints for each SV as an anchor and calculated the shortest distance along the concatenated circular genome to the other breakpoint. The majority of second breakpoints were distributed randomly throughout all chromosomes, except for a peak of intrachromosomal events less than 1Mb that were enriched more than 100 times above the Poisson mean of the random translocation artifacts (Figures 7D and E). This highly non-random portion of SV junctions identifies them as reflecting a biological rather than an error process. We considered whether the accumulation of SVs in this small size range might be an alignment artifact. However, their MAPQ values were consistently 60, the maximum reported by minimap2, and manual re-alignment using BLAT validated that both read segments flanking the junctions were properly and uniquely aligned.

**Figure 7.**
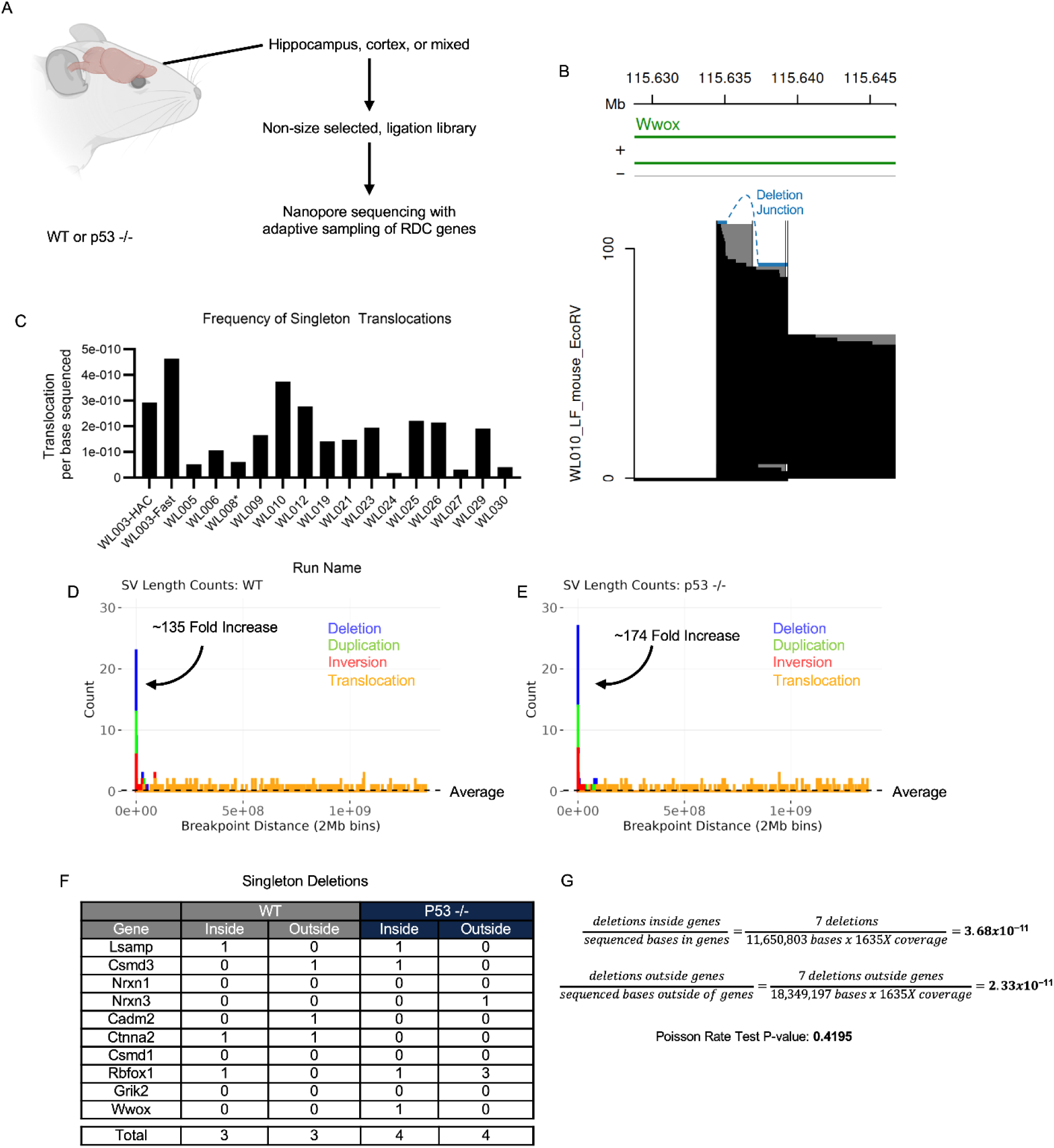
SVs in large genes in the adult mouse brain. **A.** Mouse brains were harvested from different sources and genotypes before nanopore sequencing targeting ten large RDC genes. Image created with BioRender.com. **B.** Example deletion containing read (blue) in the WWOX gene. RE sites are vertical black lines. Non-SV reads are black with projections in gray. **C.** Singleton translocations per base sequenced across all experiments. An asterisk denotes one run that was not adaptively sampled. **D. and E.** Histogram of the genomic distance the second SV breakpoint is from an anchored first breakpoint by SV type, in 2Mb bins. Dotted line depicts the average number of SVs per bin. (D) is wild-type, (E) is p53-/- mice. **F.** Singleton deletions between 10 kb and 1 Mb by gene, stratified by whether at least one breakpoint was within the corresponding gene body (inside) or in the 1 Mb padding on either side of the gene (outside). **G.** Poisson rate test calculation.

Like deletion SVs in CFSs, RDCs detected in NSPCs occur more often within gene bodies toward their centers. Although we did detect some deletions in both genotypes with at least one breakpoint inside the gene body that were between 10kb and 1Mb in size, there was no statistical significance in the frequency of deletions that occurred inside (7 events) versus outside (7 events) gene bodies after accounting for the number of bases in these regions (Figure 7F and G). While the current study is either underpowered to reach a biological conclusion about large genes or RDCs do not lead to intragenic SV accumulation in the brain, the combined results nevertheless support the accurate detection of somatic SVs by HiFiRe3.

### Bleomycin dose-dependent increase in SVs in short and long-read platforms

To further explore the ability of HiFiRe3 to detect rare biological SVs, we prepared HiFiRe3 libraries for the Aviti and PacBio platforms using GM12878 cells treated with varying doses of bleomycin (0, 1, 2.5, and 5 μM). Bleomycin is known to induce SVs by creating DSBs, especially interchromosomal events, i.e., translocations (30), unlike the intrachromosomal events expected above at large genes. We first compared the artifact frequency of libraries not treated with bleomycin across our different library approaches (Figure 8A). Aviti short read libraries had the highest frequency of singleton translocations that passed all filters per base sequenced, 6 to 19 times more than PacBio or ONT. Nevertheless, both the Aviti and PacBio platforms revealed a clear dose-dependent increase in SV formation, including singleton translocations (Figure 8B,C). These data show that HiFiRe3 can detect meaningful genotoxicant-induced SV signals above a low background noise in untargeted, reduced representation libraries.

**Figure 8.**
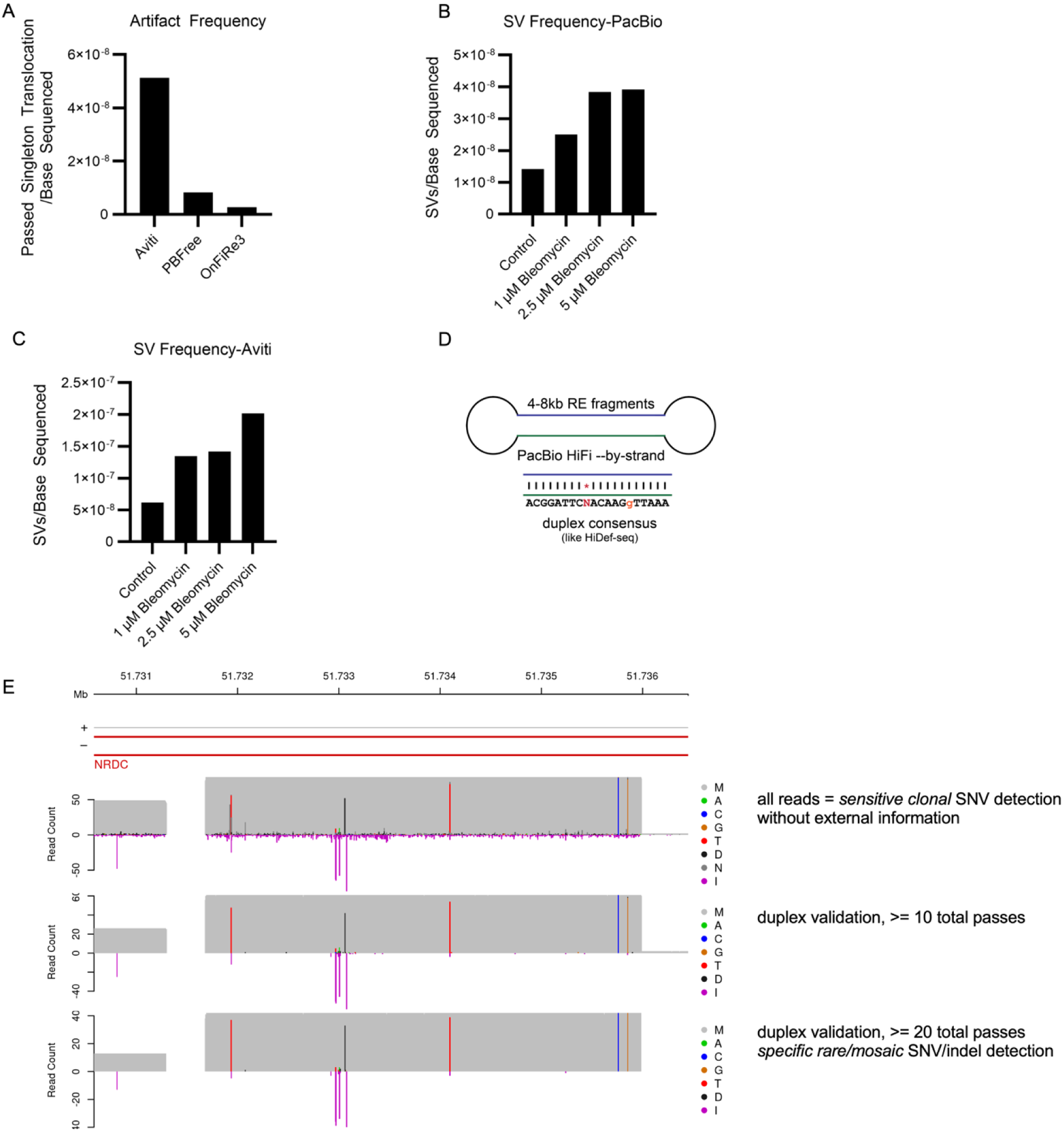
Paired detection of SVs and SNVs in the same samples using PacBio A. and. **B.** Error-corrected SVs per base sequenced at varying doses of bleomycin in Aviti (A) and PacBio libraries (B). **C.** Comparison of library artifacts rates, i.e., singleton translocation frequencies after applying error correction filters across different library types. **D.** Selecting for 4-8kb fragments and performing PacBio basecalling by strand supports duplex consensus sequence creation. **E.** Each graph depicts the number of reads matching SNVs and indels across a genome window at different filtering levels.

### SV and SNV/indel error correction in the same PacBio library

Finally, in many applications of error-corrected sequencing, tracking SNVs as well as SVs in the same library would be desirable, such as testing the effects of genotoxic agents and the accumulation of somatic variants. HiFiRe3 offers error-minimized sequencing for both SNVs and SVs using PacBio as an extension of HiDEF-seq. Like HiDEF-seq, HiFiRe3 RE fragments are size selected to a range that allows more passes per molecule during sequencing and thus construction and comparison of strand-level consensuses for duplex error correction. The key addition with HiFiRe3 are the SV error correction algorithms built into the pipeline and applied the same way as for other platforms. The same PacBio libraries discussed in the bleomycin experiment above were size-selected to 4-8kb, allowing for more than 20 passes per read (Figure 8D). In addition to detecting SVs, by applying our filters to only accept SNVs that had 20 or more passes, we could detect both clonal and specific/rare SNVs/indels within our cell populations (Figure 8E). Although bleomycin is primarily a clastogenic agent that induces structural genomic alterations, SNVs have been reported for related bleomycin-family compounds in some experimental systems (31).

## Discussion

HiFiRe3 extends error-corrected sequencing using standard adapter ligation libraries such that rare SVs can be studied with a specificity approaching SNV analyses by duplex sequencing. We anticipate applications in genetic toxicology, somatic mosaicism, and cancer and DNA repair biology. By being cognizant of both error suppression and correction, HiFiRe3 fills a niche in SV workflows, where many advances have focused on algorithmic improvements with multi-read confirmation rather than error minimization at the single-molecule level. HiFiRe3 FREE SiteMatch and <1N StemLength filters leverage two pre-ligation properties of source DNA molecules – their endpoints and sizes - to establish rules for how true SV molecules are expected to behave. Junctions that violate either expectation are enriched for abundant ONT follow-on events, ligation chimeras, and other artifact classes that can be removed without substantially reducing sensitivity for true SVs. In exploiting two simple physical properties of source DNA molecules, HiFiRe3 is platform-agnostic while still exploiting platform-specific failure modes, such as ONT follow-ons, or features, such as PacBio multi-pass reads.

Systematic analysis of SV artifacts extended the two primary error correction modes by identifying additional frequent error processes. In ONT libraries, most artifact junctions show insertions with easily identified stretches of low-quality bases and adapter sequences, hallmarks of follow-on artifacts that conjoin two physically separate DNA strands. These events include both intramolecular strand pairs appearing as inversions and intermolecular pairs appearing as translocations and widely dispersed intrachromosomal junctions. Follow-on adapter sequences are often low quality and missed by algorithms that are not junction directed or expecting internal adapters. In Aviti libraries the dominant artifact classes instead reflect blunt-end ligation and low-quality short alignments, as captured by the SiteMatch, StemLength, and MapQ filters. PacBio libraries show another pattern, with foldback inversions and lower-confidence alignments contributing most to the filtered pool.

Despite the platform-specific differences, HiFiRe3 supported a consistent interpretation of SV calls as the residual, biologically plausible subset of junctions that survived multiple independent filters across ONT, Aviti, and PacBio experiments. This convergence suggests that HiFiRe3 error logics can suppress artifactual noise and allow for bona fide DNA breakage and repair to be detected and SV rates calculated. Given the nature of the biological systems we used – replication-associated SVs in large genes and bleomycin treatment – application of the full HiFiRe3 filter set yielded junctions that had microhomology and *de novo* insertion distributions distinct from many artifact junctions but characteristic of biological end-joining. HiFiRe3 does not depend on nonhomologous junctions, however, and should reveal any SV formation mechanism consistent with the platform read length.

Biologically, HiFiRe3 with full error correction validated previous observations about SVs formed under replication stress in CFSs. In RPE-1 cells treated with APH, HiFiRe3 ONT libraries targeted to candidate loci revealed a concentration of deletions towards the centers of a subset of large genes including CDH13, GPC6, and WWOX, with a median size range consistent with reported intrachromosomal events at CFSs. From the first experimental replicate executed on one ONT flow cell in a few days, SV hotspots in RPE-1 emerged with confidence because HiFiRe3 and svCapture showed the same biases toward deletions, gene centers, and only some large genes.

At large neural genes in the mouse brain previously identified as RDCs, HiFiRe3 again performed well at detecting a much lower baseline level of singleton SVs. An aid was that artifactual SVs join random genome positions whereas many important, and the most typical, biological SVs join breakpoints separated by 10 kb to 1Mb. Indeed, breakpoint proximity alone is a valuable and relevant error correction filter for biological settings where translocations are unexpected. Despite confident SV assignments, present data do not support a statistically significant enrichment of intragenic deletions within RDC genes in vivo. However, statistical power was only around 10% with the event count obtained. A fundamental challenge when scoring locus-specific somatic SVs is that coverage must be very high to detect rare events. ONT HiFiRe3 libraries can be targeted using adaptive sampling and, as PCR-free libraries, give unique information from every read, but even so cost can become prohibitive. For mouse brain RDC genes, we estimate that 12 times more coverage than we gathered could yield a confident acceptance or rejection of the RDC-SV hypothesis. We are exploring probe capture with short-read libraries as a means of increasing coverage in target loci.

PacBio experiments showed that HiFiRe3 have the potential to jointly track SVs and SNVs in the same libraries, enabling integrated assessments of clastogenic and mutagenic outcomes. Bleomycin dose-dependent increases in singleton translocations and other SV classes were readily detected and the inferred microhomology profiles again distinguish artifact-dominated junctions from those consistent with biological end-joining. These data support the potential of HiFiRe3 for scoring biological stressors, such as a strong clastogen, that promote random chromosomal SV pairings in the same manner as artifact junctions. At the same time, per-base SNV rates could be estimated in the manner reported as HiDEF-seq, which PacBio HiFiRe3 closely resembles with the important additions of RE site matching and SV error correction.

A key property of HiFiRe3 as implemented here is that it has reduced representation, i.e., not all parts of the genome are sequenced when using RE fragments. This is often a valued feature as HiFiRe3 piles up reads to allow identification of clonal SVs from the subject library without external comparators, especially when using strict size selection and region targeting with ONT adaptive sampling. Notably, despite being reduced representation, HiFiRe3 can reveal *de novo* SV junctions throughout the entirety of the genome or of a targeted region. It simply reveals the subset of those junctions that pass RE fragment sizing. However, at times it is desirable to capture all possible SV junctions, which can be achieved by applying size selection to sheared genomic DNA molecules before adapter ligation to enable StemLength while foregoing SiteMatch filtering. We do not report such libraries here but demonstrate the redundance of these and other filters due to shared properties of SV artifact molecules, such that we believe SV error rates could be greatly improved in true whole-genome sequencing, including genome assembly where even rare SV errors in libraries can be confounding. This ability to adapt to common existing library protocols was a goal in HiFiRe3 design.

## Methods

### Cell lines and growth

#### GM12878 lymphoblastoid cells

GM12878, (Coriell, RRID CVCL_7526), is a XX female, euploid, EBV-immortalized human lymphoblastoid cell line generated from the HapMap Project. GM12878 cells were cultured at 37C with 5% CO2 in RPMI 1640 medium supplemented with 15% FBS, 2mM L-glutamine, and 100 U/mL penicillin-streptomycin.

#### RPE-1 epithelial cells

RPE-1, (hTERT RPE-1), is an immortalized human female retinal pigment epithelial cell line. These cells were gifted by the lab of Jayakrishnan Nandakumar (University of Michigan), validated by STR profiling (ATCC) and tested for mycoplasma. RPE-1 cells were cultured at 37C with 5% CO12 in DMEM/F12 medium supplemented with 10% FBS, 2mM L-glutamine, and 100 U/mL penicillin-streptomycin.

#### Cell treatment with bleomycin and aphidicolin

When indicated for SNV and SV induction experiments, cells were treated with varying concentrations of bleomycin. Bleomycin (Cayman Chemical Company) was dissolved in DMSO at a stock concentration of 10 mM. GM12878 cells were cultured at 1 μM, 2.5 μM, and 5 μM bleomycin for 24 hours and then allowed to recover in fresh media for 72 hours before harvesting.

When indicated for SV induction experiments, cells were treated with the DNA polymerase inhibitor aphidicolin (APH). APH (Sigma) was dissolved in DMSO at a stock concentration of 200 μM. GM12878 cells were treated with 0.4 μM APH for 48 hours before harvesting. RPE-1 cells were treated with 0.6 μM APH for 72 hours before harvesting.

#### Mouse brain sources and handling

Three total wild-type mouse brain samples were supplied by the Schwer lab (University of California San Francisco) while two total p53^-/-^ mouse brain samples were supplied by the Sekiguchi lab (University of Michigan) and Jackson laboratories (002101 - B6.129S2-Trp53<TM1TYJ>/J - HOM -). Samples from the Schwer lab and Jackson laboratories were dissected before delivery into cortex and hippocampus tissue. Sekiguchi lab tissue was dissected by the Wilson lab to harvest cortex tissue. Detailed information about individual mouse age, tissue, and sequencing runs can be found in Supplemental Table 1.

#### Summary of reported libraries

A total of 29 HiFiRe3 libraries in different combinations of library preparation strategy and sequencing platform are reported in this study, as tabulated in Supplemental Table 2. Sequencing files as obtained from the platform via the sequencing facility are available as described in Data Availability.

#### DNA extraction, fragmentation, and library preparation

All libraries reported in this study used an approach in which DNA extraction and fragmentation were combined in a concerted process using one of the New England Biolabs (NEB) Monarch high molecular weight (HMW) genomic DNA extraction kits where DNA was released from the extraction beads by digestion with a restriction enzyme (RE) before further processing.

##### DNA extraction onto beads

We extracted genomic DNA using either the Monarch HMW DNA Extraction Kit for Tissue (T3060L) or the Monarch HMW DNA Extraction Kit for Cells & Blood (T3050L). The input amount for cell lines varied from 2-6 million cells. Cells were washed with 1X PBS before pelleting by centrifugation at 700 x g for 5 minutes at 4°C. The input amount for mouse brain tissue was 15-20 mg. We incubated the cells or tissue in lysis buffer and Proteinase K at 37°C with shaking at 250 rpm and followed the NEB protocols for DNA adherence to beads and bead washing.

We found that we could also use the tissue kit for cells. We modified the DNA extraction protocol published by Oxford Nanopore Technologies (ONT) for the Ultra-Long DNA Sequencing Kit V14 (SQK-ULK114) which uses the Monarch HMW DNA Extraction Kit for Tissue. We incubated the cells with lysis buffer and Proteinase K at 37°C for 30 minutes in a thermoblock with shaking at 250 rpm. The RNase digestion ran for 10 minutes at 37°C but with no shaking. We did not perform the protein separation step when extracting DNA from cells with this method.

##### DNA release via restriction enzyme fragmentation

For all extraction methods, we pulse spun to remove excess gDNA wash buffer and poured the final washed beads with bound DNA into a 2 ml round bottom tube with a RE cocktail containing 100 to 400 units depending on cell count and 1X CutSmart Buffer in an appropriately scaled volume of 100 to 400 µL. Digestion proceeded at 37°C for one hour for AluI and EcoRV-HF enzymes. To distribute the enzyme cocktail around the beads, we gently rolled and ½ inverted the tubes every 10 minutes. After one hour, we poured the beads and solution into the Monarch Bead Retainer in a 1.5 ml tube and separated the released DNA from the beads by pulse spin (short reads) or by gravity (long reads).

##### A-tailing and nick blocking

After RE digestion, we quantified the DNA using the Qubit dsDNA BR (Broad Range) Assay Kit using a volume of 10 µL of digested DNA. For initial libraries, without size selection by Blue Pippin, we diluted 2 µg of DNA into a 100 µL total volume solution of water and 1X CutSmart buffer, 30 units Klenow exo- and 2.5 mM dATP/ddBTP. We scaled up the A-tailing and nick blocking reaction for DNA destined for size selection by the Blue Pippin based on quantification and the maximum capacity of 5 μg per lane. To avoid mechanical shearing, we used a wide-bore pipette to gently mix the A-tailing solution 3-4 times. We incubated the A-tailing solution at 37°C in a thermoblock with no shaking.

##### Bead cleanup

For non-Pippin size-selected libraries, we purified the A-tailed and blocked DNA using AMPure beads at a 0.8X bead to DNA ratio. Alu-digested libraries used a 1.6X bead to DNA ratio. We used a 0.8X bead cleanup ratio for EcoRV-HF-digested libraries. We eluted DNA off the magnetic beads with either nuclease-free water, 1X TE, or low EDTA TE. Final elution volumes varied according to the library preparation method used for the target sequencing platform.

##### Size selection with the BluePippin

AluI digested libraries for Aviti were size selected using the BLF2003 cassette and the 2% DF 100-600 bp marker M1 cassette definition (Sage Science). For PacBio and ONT libraries we used the BLF7510 cassette and the 0.75% DF 3–10kb Marker S1 – Improved Recovery cassette definition (Sage Science). The PacBio libraries used a 4600 – 8000 bp range collection. The ONT libraries used 8050 to 16000 bp range collection. We allowed the DNA in the elution well to rest for 3-8 hours. To improve recovery from the elution well we added 5-10 µL of 0.1% Tween 20 in electrophoresis buffer (supplied with the cassettes) and allowed it to equilibrate in the well for 30 minutes. To avoid mechanical shearing, we used a large orifice, non-binding 300 µL tip which was pre-wet with the same Tween solution above and aspirated the DNA by slowly turning the thumbwheel on the pipette. The Pippin buffer is compatible with all library preparation methods.

##### TruSeq adapter ligation

We used the TruSeq DNA PCR-Free Library Prep Kit from Illumina (20016327) for Aviti libraries. TruSeq adapter ligation requires the ATL buffer. Because our DNAs were already blunt-ended and A-tailed, we inactivated the enzymes in the ATL buffer by incubating the ATL solution at 70°C for 5 minutes in a thermocycler. We proceeded with TruSeq adapter ligation using our size-selected DNA and the cooled ATL buffer after the A-tailing step of the protocol. We used Unique Dual Indices (UDIs) from TruSeq (IDT for Illumina TrueSeq DNA UD Indexes v2).

##### ONT adapter ligation

DNA from the BluePippin went directly into the ONT Ligation Kit protocol. In the case of size selected libraries, it is important to use a 1:1 bead clean-up ratio after ligation. For all long-read libraries, we used the Long Fragment Buffer.

##### PacBio adapter ligation

PacBio library prep began at the ABC step by adding 8 µL of Repair Buffer to the DNA from the Pippin and bringing the total volume to 60 µL with low EDTA TE. From there, we followed the manufacturer’s instructions for the SMRTBELL 3.0 Library Prep Kit (102-141-700) and used SMRTBELL adapter index plate A (102-009-200)

#### High throughput sequencing

ONT sequencing was performed on a PromethION P2 sequencer in the Wilson laboratory using v14 chemistry and PromethION flowcells (FLO-PRO114M). When indicated, adaptive sampling was performed using a BED file of regions corresponding to approximately 1% of the genome using Fast base calling on the P2 device.

All other sequencing was performed at the University of Michigan Advanced Genomics Core on either an ONT PromethION P24 using v14 chemistry and PromethION flowcells (FLO-PRO114M), a PacBio Revio, or an Aviti at Element Bioscience’s Service Lab using FreeStyle chemistry. Aviti libraries were sequenced as either paired-end 2x150 or single-end 1x300 base pairs. All Aviti run recipes started with 2 bases of dark cycling to account for the expected two fixed 5’ bases resulting from digestion with AluI.

#### Data analysis in the HiFiRe3 tool suite

The HiFiRe3 tool suite was assembled in the flow control and repository system of the Michigan Data Interface (MDI, https://midataint.github.io) and is publicly available on GitHub (https://github.com/wilsontelab/HiFiRe3). It carries data analysis pipelines accessed through a Linux command line interface (CLI) and R Shiny visualization apps runnable on any platform. Computation tools include third-party programs mostly installed into a Conda environment, which is available in a version-controlled Singularity Apptainer or can be built locally. HiFiRe3-specific code is mostly written in Rust, with the pre-compiled hf3_tools binary available via GitHub releases. The container and binary are downloaded automatically by the CLI. Documentation and configuration examples are provided in the repository. In best usage, the library type, sequencing platform, restriction enzyme, size selection, reference genome, and other sample parameters are described in a job configuration file. The CLI and binary encompass the full range of supported platforms and library types, which are processed through platform-specific and shared pipeline actions described below.

#### Basecalling

Two sequencing platforms require specific HiFiRe3 basecalling. ONT POD5 raw data files are basecalled using ‘dorado’ (https://github.com/nanoporetech/dorado) in pipeline action ‘basecall ONT’. where all samples in this study used the highest accuracy ‘sup’ model. Dorado runs without adapter trimming (option ‘--no-trim’) so that trimming can be performed in a RE-aware manner before writing unaligned BAM files. For example, a six-base blunt EcoRV site, 5’-GAT|ATC, puts bases ATC at the 5’ end of all cut fragments. Module ‘trim_ont’ uses a fast Smith-Waterman search at read ends for the ONT adapter plus these three fixed bases for more accurate adapter localization than with trimmers unaware of this configuration. Trimming outcome is reported in BAM ‘tl:Z:’ tag along with retained Dorado channel and other tags.

HiFiRe3 libraries of type PacBioMolecule are basecalled on Revio or Vega in the normal mode of two-strand consensus calling. For error-corrected SNV calling, HiFiRe3 PacBioStrand libraries are basecalled on the sequencer using option ‘--by-strand’ to obtain a consensus for each strand individually. Pipeline action ‘basecall PacBio’ scans the unaligned BAM files for strand pairs which are then aligned to each other and to the reference genome to support three-strand error correction. Base differences along the strands are encoded into tags ‘dt:i:’ and ‘dd:Z:’ (see Rust code for format details) along with other retained PacBio tags. Homoduplex bases where the two sequenced strands agree are reported in the final read sequence regardless of their reference match. Heteroduplex strands where one base span matches the reference are reported as the reference bases with kinetic information at the mismatched bases put into tag ‘sk:B:S,’. Positions where all three strands differ are reported as one more N bases to prevent them from calling SNVs downstream.

#### Read alignment

All library types are aligned to the relevant reference genome using ‘minimap2’ (32), including short reads, in pipeline action ‘analyze fragments’ with options ‘-y -Y --secondary=no’. Short-read libraries are prepared for alignment by ‘fastp’ (33) to perform any required adapter trimming, initial quality filtering, and merging of overlapping read pairs on paired end platforms (tag ‘fm:Z:’). For multi-species libraries, we prepared a concatenated reference genome of all chromosomes from both species with chromosome names suffixed with the UCSC genome name, e.g., chr1_hg38, using pipeline action ‘combine genomes’. HiFiRe3 tracks reads by genome with this naming convention if option ‘--is-composite-genome’ is set. Bandwidth during alignment is set to ‘100’ for short reads and ‘500,3300’ for long reads to encourage the aligner to split reads with SV junctions on the scale of the read length into primary and supplementary alignments rather than encoding them in the CIGAR string and ‘cs:Z:’ tags of a single alignment.

#### Initial readh3 and alignment metadata

Initial analysis of sequencing reads is performed immediately following alignment before writing name-sorted BAM files. Module ‘analyze_alignments’ in hf3_tools performs tasks that do not require knowledge about RE sites or insert sizes. Unmerged paired-end reads are split and considered as two independent reads because the unsequenced bases in the read center preclude accurate assessment of insert sizes and are untrustworthy with respect to their potential to include chimeric junctions. All alignments of each read are ordered by their 5’ position along the read. Tags describe whether the 5’ end of the alignment matched a target region (tags ‘tm:i’. ‘to:A:’), fell below a settable MAPQ threshold, had a high aligned base divergence, were excessively short, or had low average base quality, as packed into alignment failure tag ‘af:i:’ as flag bits Mapq(1), Divergence(2), FlankLen(4), and BaseQual(8), respectively.

Read paths are characterized as read outer endpoints (tag ‘on:Z:’) and any SV breakpoint nodes, i.e., the closest reference bases in the alignments flanking each side of the junction. Breakpoint nodes are used to calculate junction attributes packed into junction tag ‘jx:Z:’. Junction types, i.e., deletion, duplication, inversion, and translocation, are determined in standard fashion from the placement and orientation of flanking alignments (34). Alignment offset is the extent to which breakpoint node positions overlap in the read sequence. When flanking alignments share overlapping read bases the inferred microhomology is calculated as a negative alignment offset, blunt junctions have a zero offset, and inserted bases not accounted for by any reference bases lead to a positive offset.

Final metadata at this stage are packed into initial junction failure tag ‘ji:i:’ Traversal data calculates the difference in the number of read bases traversed on the read and on a single chromosome between all pairs of breakpoint nodes in a set of SV read alignments. When any two nodes have a traversal delta smaller than a threshold (50 bp for short reads, 1kb for long reads), they are likely to be artifacts created by low quality base stretches that could not be aligned properly. This leads to either false deletions with an approximately equal number of apparently inserted bases or incorrect alignments to a distant genomic location with two adjacent artifact junctions. All junctions between two nodes that fail traversal delta (flag bit Traversal(1)) are numbered as part of the same reference block (tag ‘bn:i:’). The initial chimeric read splitter finally flags junctions at foldback inversions, i.e., alignment pairs that are overlapping reverse complements at a genomic locus (flag bit FoldbackInv(4)), have larger insertions with stretches of low quality bases (flag bit LowQualIns(8), or where a sequencing adapter could be aligned to any *de novo* junction base insertions (flag bit HasAdapter(16).

#### In silico RE digestion and polymorphic site finding

SV error correction by forced restriction enzyme ends (FREE) depends on knowing which genomic positions represent RE sites and are thus expected at fragment ends but not in their middles. HiFiRe3 finds genomic RE site positions first by efficient *in silico* digestion of the reference genome in pipeline action ‘prepare genome’ to create lists of RE site positions identified as the first base after the cleaved phosphodiester bond on the top genome strand. However, because not all RE sites in a DNA sample will match the reference due to polymorphisms, outer endpoint nodes from the initial alignment analysis are tabulated to identify sites that appear at read ends with a frequency comparable to *in silico* sites. The position of the outermost aligned read base is adjusted as needed to account for any clipped bases as well as the alignment strand to match the site position as defined above. Polymorphic plus *in silico* sites are collected as ‘filtering sites’ for use in RE site matching below.

#### RE site-aware read metadata

Armed with sample-specific filtering site positions, hf3_tools module ‘analyze_inserts’ continues to tag reads and alignments with error correction metadata. FREE error correction adds failure flag SiteMatch(32) to the final junction failure tag ‘jf:i:’ when either breakpoint node fell within a threshold distance from a filtering site (25bp for ONT, 10bp for PacBio, 5bp for short read platforms), with tag ‘sc:B:i,’ carrying the distances. Such ends are likely to represent end-to-end chimeras. Entire reads are also flagged in read failure tag ‘rf:i:’ if their outer endpoints do not match expected RE sites, where different sequencing platforms sequence different parts of the original DNA inserts. PacBio and paired-end short read platforms are obligatorily end-to-end, i.e., the read outer endpoints reliably correspond to the insert endpoints. In contrast, many reads on single-end platforms such as Aviti only report a true insert outer endpoint at the read 5’ end when the read 3’ end falls short of the second outer endpoint in a long insert. ONT reads often extend to the 3’ outer endpoint to create an end-to-end read, as confirmed by productive adapter trimming at that end, but sometimes truncate before reaching the insert 3’ end. When reads are not end-to-end, hf3_tools projects the observed read 3’ end to the nearest RE filtering site as a prediction of the RE fragment that likely gave rise to it.

#### Size selection-derived read metadata

The 5’ and projected 3’ outer endpoints are used to infer DNA insert sizes in RE-digested libraries because they account for any unsequenced 3’ bases. Initially, options ‘--min-selected-size’ and ‘--selected-size-cv’ communicate the expected lower limit of selected fragments and the precision of that boundary, which the pipeline uses to calculate the 1N and 2N limit values. Sometimes examination of inferred insert size distributions allows those values to be refined by setting option ‘--min-allowed-size’ as a forced value for 1N. Size-based error correction metadata is then populated by calculating the stem length of any junction, defined as the shortest distance from the junction to either validated (not projected) outer endpoint of the read (tags ‘ul:i:’ and ‘vl:i:’). Any read with a stem length >1N has flag bit StemLength(64) set in jxn failure tag ‘jf:i:’ as suspicious for being an end-to-end chimera.

Notably, initial junction failure flags Traversal, FoldbackInv, LowQualIns and HasAdapter are applied exclusively in that order such that no later flag will be set if an earlier flag failed. In contrast, SiteMatch and StemLength flags are both set when appropriate for a library type, or otherwise are left unset to always pass that filter.

#### SV finding, aggregation, and error correction

SV analysis next entails fuzzy grouping of individual annotated read junctions into “final junctions” thought to represent independent observations of the same SV. First, breakpoint nodes are reordered into a canonical orientation by sorting them by their position along the concatenated reference genome. Ordered junctions are sorted and compared to identify clusters of 5’ breakpoint nodes followed by nested clusters of 3’ breakpoint nodes within the 5’ groups, where a node cluster falls within 500bp on ONT due to its lower base accuracy and 20bp on other platforms.

We next account for alignment clipping due to base mismatches, where inappropriately clipped junction bases do two things to the same degree: (i) decrease the distance from the breakpoint node to the chromosome end, called the chromosome stem length, and (ii) increase the alignment offset, i.e., the number of apparently inserted bases. Thus, the final clustering metric is the summed chromosome stem lengths on the two sides of the junction plus the alignment offset, which is an SV rather than a breakpoint-level assessment resilient to errors on either or both sides of the junction. It applies to all types of junctions, including inversions and translocations, regardless of whether additional junctions exist in the read.

Fuzzy-matching yields final junctions, each with a set of one or more matching read instances. The most frequently observed node pair in each set is used to describe the junction. Matching reads are unique for most platforms and library types with two exceptions. First, ONT can sequence two strands from the same molecule as independent reads in the same pore, so each nanopore channel is only allowed to count reads on one DNA strand for a given junction (reads from the same strand in one channel must represent different source molecules). HiFiRe3 also handles PCR-amplified short reads by deduplicating junction instances based on shared read outer endpoints. The outcome is a final junctions table (see Rust code for its format).

Manipulations above can be applied to a single sample by pipeline action ‘analyze SVs’ or to the combined data from samples of the same library type by ‘compare SVs’. Analyzing combined samples affords the best fuzzy matching, but libraries of different types can also be compared after initial analysis using ‘merge SVs’.

Importantly, no filtering, i.e., post-alignment removal of reads or junctions, is applied through any of the prior processing. All data persist with metadata to support error correction and variant calling, e.g., by requiring multiple unique observations to call clonal junctions or applying error correction filters to singleton junctions to call rare mosaic SVs. For this manuscript, filters were applied in the R Shiny app, which also constructed plots of junction properties.

#### SNV finding, aggregation, and error correction

SNV calling is restricted to PacBioStrand libraries that support duplex error correction. It occurs in two passes over individual alignments. The first pass includes all reads, even non-duplex reads, for maximal sensitivity for clonal variant calling, given that some genome positions have lower coverage in a reduced representation library. The second pass includes only duplex reads for maximal specificity for rare mosaic variant calling. In each pass, the ‘dd:Z:’ tag (see Basecalling) is used to create a mask that dictates which bases of a read are allowed to call error-corrected SNVs and indels. The minimap2 ‘cs:Z:’ tag is then scanned to construct a list of unique variants and a pileup of observed bases at each genome position. Each record type includes the number of observations that were allowed or disallowed by the homoduplex mask. Comments above for SVs regarding sample comparison vs. merging, filtering of variants calls by exploiting metadata, and visualization in the R Shiny app apply equally to SNVs.

## Acknowledgements

Funding for this work was provided by NIH grants R21 ES022311 to T.E.W. and R01 GM147026 to T.E.W. and T.W.G. We thank Sarah Cohen and Pavlina Chuntova for harvesting brain tissue. We thank the Sekiguchi lab at the University of Michigan for the donation of mouse tissue as well as use of their tissue culture facilities. We thank Li Ching Chen for assistance examining mixed species libraries and other HiFiRe3 variants not reported here.

## Author Contributions

T.E.W. administered and conceived the study. All authors were involved in sample collection, data collection, and/or data analysis. J.A.S. and T.E.W. wrote the manuscript.

## Code Availability

The HiFiRe3 data analysis tool set is publicly available on GitHub at https://github.com/wilsontelab/HiFiRe3.

## Data Availability

Basecalled reads for all reported samples will be made available at the Sequence Read Archive upon publication.

## References

1. Kaivola, K., Chia, R., Ding, J., Rasheed, M., Fujita, M., Menon, V., Walton, R.L., Collins, R.L., Billingsley, K., Brand, H. et al. (2023) Genome-wide structural variant analysis identifies risk loci for non-Alzheimer&#x27;s dementias. Cell Genom, 3, 100316.

2. Zhou, B., Arthur, J.G., Guo, H., Kim, T., Huang, Y., Pattni, R., Wang, T., Kundu, S., Luo, J.X.J., Lee, H. et al. (2024) Detection and analysis of complex structural variation in human genomes across populations and in brains of donors with psychiatric disorders. Cell, 187, 6687–6706.e6625.

3. Gillani, R., Collins, R.L., Crowdis, J., Garza, A., Jones, J.K., Walker, M., Sanchis-Juan, A., Whelan, C.W., Pierce-Hoffman, E., Talkowski, M.E. et al. (2025) Rare germline structural variants increase risk for pediatric solid tumors. Science, 387, eadq0071.

4. Kennedy, S.R., Schmitt, M.W., Fox, E.J., Kohrn, B.F., Salk, J.J., Ahn, E.H., Prindle, M.J., Kuong, K.J., Shen, J.C., Risques, R.A. et al. (2014) Detecting ultralow-frequency mutations by Duplex Sequencing. Nat Protoc, 9, 2586–2606.

5. Valentine, C.C., 3rd, Young, R.R., Fielden, M.R., Kulkarni, R., Williams, L.N., Li, T., Minocherhomji, S. and Salk, J.J. (2020) Direct quantification of in vivo mutagenesis and carcinogenesis using duplex sequencing. Proc Natl Acad Sci U S A, 117, 33414–33425.

6. Bhawsinghka, N., Burkholder, A. and Schaaper, R.M. (2023) Detection of DNA replication errors and 8-oxo-dGTP-mediated mutations in E. coli by Duplex DNA Sequencing. DNA Repair (Amst), 123, 103462.

7. Pilheden, M., Ahlgren, L., Hyrenius-Wittsten, A., Gonzalez-Pena, V., Sturesson, H., Hansen Marquart, H.V., Lausen, B., Castor, A., Pronk, C.J., Barbany, G. et al. (2022) Duplex Sequencing Uncovers Recurrent Low-frequency Cancer-associated Mutations in Infant and Childhood KMT2A-rearranged Acute Leukemia. Hemasphere, 6, e785.

8. Xiong, K., Shea, D., Rhoades, J., Blewett, T., Liu, R., Bae, J.H., Nguyen, E., Makrigiorgos, G.M., Golub, T.R. and Adalsteinsson, V.A. (2022) Duplex-Repair enables highly accurate sequencing, despite DNA damage. Nucleic Acids Res, 50, e1.

9. Schmitt, M.W., Kennedy, S.R., Salk, J.J., Fox, E.J., Hiatt, J.B. and Loeb, L.A. (2012) Detection of ultra-rare mutations by next-generation sequencing. Proc Natl Acad Sci U S A, 109, 14508–14513.

10. Bae, J.H., Liu, R., Roberts, E., Nguyen, E., Tabrizi, S., Rhoades, J., Blewett, T., Xiong, K., Gydush, G., Shea, D. et al. (2023) Single duplex DNA sequencing with CODEC detects mutations with high sensitivity. Nat Genet, 55, 871–879.

11. Maslov, A.Y., Makhortov, S., Sun, S., Heid, J., Dong, X., Lee, M. and Vijg, J. (2022) Single-molecule, quantitative detection of low-abundance somatic mutations by high-throughput sequencing. Sci Adv, 8, eabm3259.

12. Cheng, A.P., Rusinek, I., Sossin, A., Widman, A.J., Meiri, E., Krieger, G., Hirschberg, O., Tov, D.S., Gilad, S., Jaimovich, A., et al. (2025) Paired plus-minus sequencing is an ultra-high throughput and accurate method for dual strand sequencing of DNA molecules. bioRxiv.

13. Lawson, A.R.J., Abascal, F., Nicola, P.A., Lensing, S.V., Roberts, A.L., Kalantzis, G., Baez-Ortega, A., Brzozowska, N., El-Sayed Moustafa, J.S., Vaitkute, D. et al. (2025) Somatic mutation and selection at population scale. Nature, 647, 411–420.

14. Liu, M.H., Costa, B.M., Bianchini, E.C., Choi, U., Bandler, R.C., Lassen, E., Grońska-Pęski, M., Schwing, A., Murphy, Z.R., Rosenkjær, D. et al. (2024) DNA mismatch and damage patterns revealed by single-molecule sequencing. Nature, 630, 752–761.

15. Abascal, F., Harvey, L.M.R., Mitchell, E., Lawson, A.R.J., Lensing, S.V., Ellis, P., Russell, A.J.C., Alcantara, R.E., Baez-Ortega, A., Wang, Y. et al. (2021) Somatic mutation landscapes at single-molecule resolution. Nature, 593, 405–410.

16. Wilson, T.E., Ahmed, S., Higgins, J., Salk, J.J. and Glover, T.W. (2023) svCapture: efficient and specific detection of very low frequency structural variant junctions by error-minimized capture sequencing. NAR Genom Bioinform, 5, lqad042.

17. Hagiwara, A., Yamatani, I., Kudoh, R., Hiramatsu, K., Kadota, J.I. and Komiya, K. (2025) Association between the Cardiothoracic Ratio on Chest X-rays and the Respiratory Function in Patients with Interstitial Lung Diseases: a Cross-sectional Study. Intern Med, 64, 1025–1030.

18. Jiang, T., Liu, Y., Jiang, Y., Li, J., Gao, Y., Cui, Z., Liu, Y., Liu, B. and Wang, Y. (2020) Long-read-based human genomic structural variation detection with cuteSV. Genome Biol, 21, 189.

19. Heller, D. and Vingron, M. (2019) SVIM: structural variant identification using mapped long reads. Bioinformatics, 35, 2907–2915.

20. Smolka, M., Paulin, L.F., Grochowski, C.M., Horner, D.W., Mahmoud, M., Behera, S., Kalef-Ezra, E., Gandhi, M., Hong, K., Pehlivan, D. et al. (2024) Detection of mosaic and population-level structural variants with Sniffles2. Nat Biotechnol, 42, 1571–1580.

21. Collins, R.L., Brand, H., Karczewski, K.J., Zhao, X., Alföldi, J., Francioli, L.C., Khera, A.V., Lowther, C., Gauthier, L.D., Wang, H. et al. (2020) A structural variation reference for medical and population genetics. Nature, 581, 444–451.

22. Chen, X., Schulz-Trieglaff, O., Shaw, R., Barnes, B., Schlesinger, F., Källberg, M., Cox, A.J., Kruglyak, S. and Saunders, C.T. (2016) Manta: rapid detection of structural variants and indels for germline and cancer sequencing applications. Bioinformatics, 32, 1220–1222.

23. Cameron, D.L., Schröder, J., Penington, J.S., Do, H., Molania, R., Dobrovic, A., Speed, T.P. and Papenfuss, A.T. (2017) GRIDSS: sensitive and specific genomic rearrangement detection using positional de Bruijn graph assembly. Genome Res, 27, 2050–2060.

24. Wilson, T.E., Ahmed, S., Winningham, A. and Glover, T.W. (2024) Replication stress induces POLQ-mediated structural variant formation throughout common fragile sites after entry into mitosis. Nat Commun, 15, 9582.

25. Liao, W.W., Asri, M., Ebler, J., Doerr, D., Haukness, M., Hickey, G., Lu, S., Lucas, J.K., Monlong, J., Abel, H.J. et al. (2023) A draft human pangenome reference. Nature, 617, 312–324.

26. Shimotsu, H. and Henner, D.J. (1984) Characterization of the Bacillus subtilis tryptophan promoter region. Proc Natl Acad Sci U S A, 81, 6315–6319.

27. Wilson, T.E., Arlt, M.F., Park, S.H., Rajendran, S., Paulsen, M., Ljungman, M. and Glover, T.W. (2015) Large transcription units unify copy number variants and common fragile sites arising under replication stress. Genome Res, 25, 189–200.

28. Wei, P.C., Chang, A.N., Kao, J., Du, Z., Meyers, R.M., Alt, F.W. and Schwer, B. (2016) Long Neural Genes Harbor Recurrent DNA Break Clusters in Neural Stem/Progenitor Cells. Cell, 164, 644–655.

29. Cabelli, V.J. and Pickett, M.J. (1953) The significance of lactose fermentation and its relationship to resistance in Klebsiella pneumoniae. J Bacteriol, 66, 443–447.

30. Bi, D., Li, X., Milić, J.V., Kubicki, D.J., Pellet, N., Luo, J., LaGrange, T., Mettraux, P., Emsley, L., Zakeeruddin, S.M. et al. (2018) Multifunctional molecular modulators for perovskite solar cells with over 20% efficiency and high operational stability. Nat Commun, 9, 4482.

31. Yu, Y., Inamdar, K.V., Turner, K., Jackson-Cook, C.K. and Povirk, L.F. (2002) Base substitutions, targeted single-base deletions, and chromosomal translocations induced by bleomycin in plateau-phase mammary epithelial cells. Radiat Res, 158, 327–338.

32. Li, H. (2021) New strategies to improve minimap2 alignment accuracy. Bioinformatics, 37, 4572–4574.

33. Chen, S., Zhou, Y., Chen, Y. and Gu, J. (2018) fastp: an ultra-fast all-in-one FASTQ preprocessor. Bioinformatics, 34, i884–i890.

34. Laufer, V.A., Glover, T.W. and Wilson, T.E. (2023) Applications of advanced technologies for detecting genomic structural variation. Mutat Res Rev Mutat Res, 792, 108475.

