## Supplemental Figures for "High-fidelity rare structural variant detection with HiFiRE3 reduced representation via restriction enzyme ends"

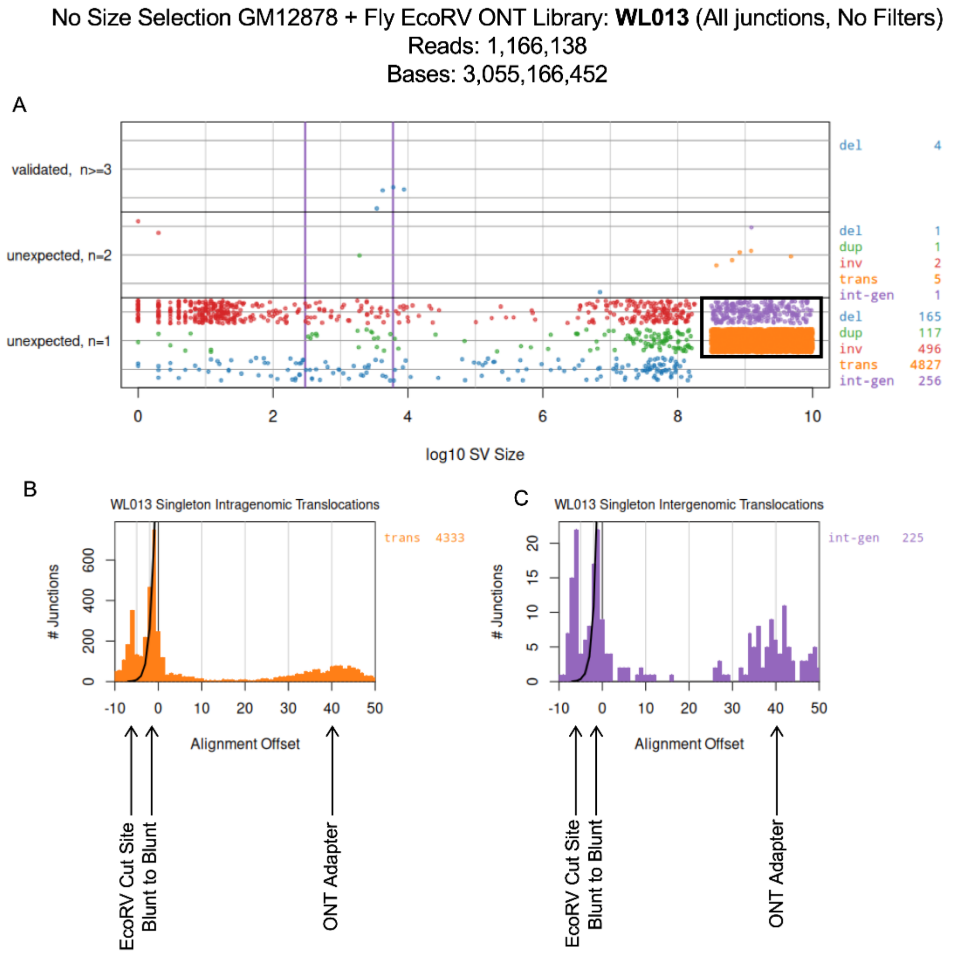


**Figure S1. Analysis of interspecies artifacts**

**A.** Junction summary plot like Figure 4A for a non-size-selected human and fly ONT library without any active filters. The black box depicts read subsets used in panels B and C.

**B. and C.** Alignment offset plot for singleton intragenomic translocations (B) and singleton intergenomic translocations (C).

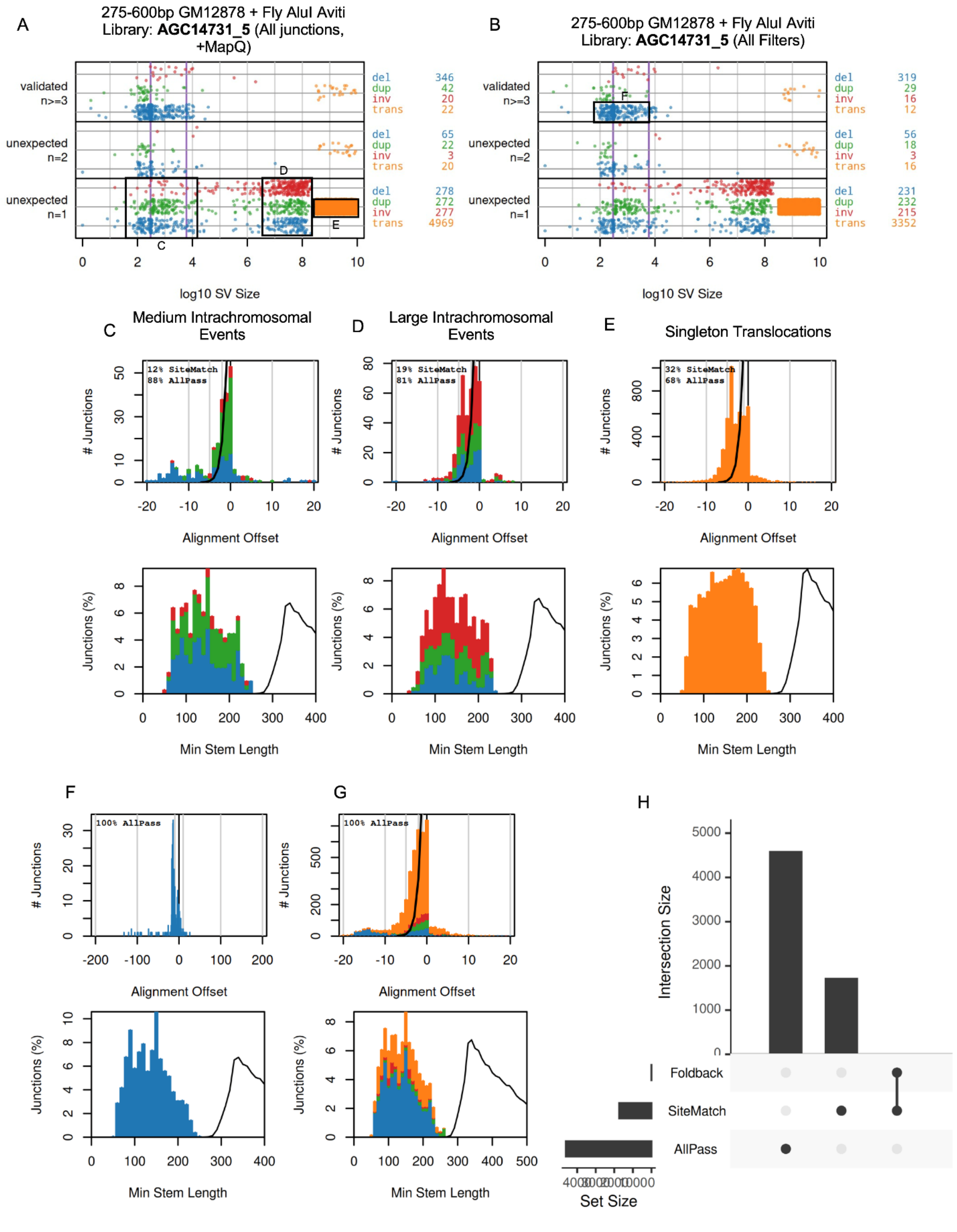
A
275-600bp GM12878 + Fly AluI Aviti
Library: AGC14731_5 (All junctions, +MapQ)
validated n>=3
unexpected n=2
unexpected n=1
del 346
dup 42
inv 20
trans 22
del 65
dup 22
inv 3
trans 20
del 278
dup 272
inv 277
trans 4969
log10 SV Size
B
275-600bp GM12878 + Fly AluI Aviti
Library: AGC14731_5 (All Filters)
del 319
dup 29
inv 16
trans 12
del 56
dup 18
inv 3
trans 16
del 231
dup 232
inv 215
trans 3352
C
Medium Intrachromosomal Events
12% SiteMatch
88% AllPass
D
Large Intrachromosomal Events
19% SiteMatch
81% AllPass
E
Singleton Translocations
32% SiteMatch
68% AllPass
### Junctions
Alignment Offset
Junctions (%)
Min Stem Length
F
100% AllPass
G
100% AllPass
H
Intersection Size
Foldback
SiteMatch
AllPass
Set Size

**Figure S2. AGC14731_5 Aviti library complete artifact analysis with MapQ filter applied**

**A.** Junction summary plot like Figure 4A for a non-size-selected human and fly Aviti 1x300 library with only the MapQ filter applied. Black boxes depict read subsets used in panels C through E.

**B.** Like A, but with all filters active. The black box depicts the read subset used in panel F.

**C. to E.** Histograms showing the number of junctions with insertions (positive alignment offset) or microhomology (negative alignment offset) (top) and minimum stem lengths (bottom) for medium intrachromosomal events, large intrachromosomal events, and singleton translocations. Insets show junction failure percentages by filter type.

**F.** Histograms like C through E now showing filtered, validated deletions.

**G.** Alignment offset plot (top) and minimum stem length plot (bottom) of remaining junctions when all filters are applied.

**H.** Upset plot showing the relationship between different filtering flags for all library junctions.

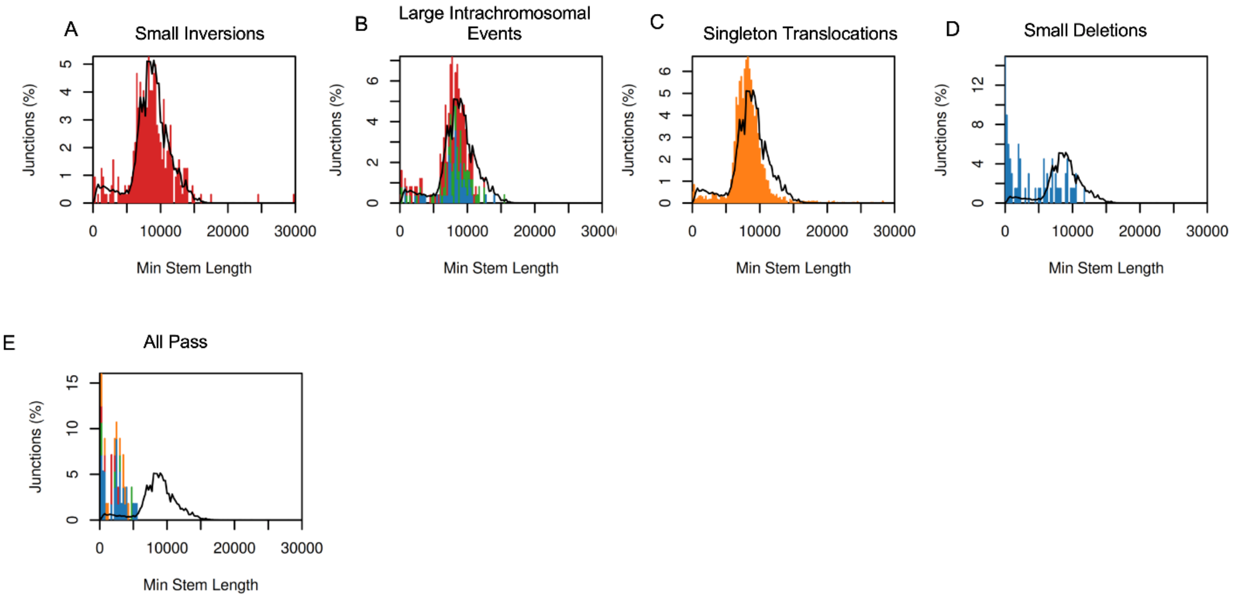


**Figure S3. WL054 stem length distributions**

**A. to E.** Minimum stem length histograms for boxed areas in Figure 4A, small inversions, large intrachromosomal events, singleton translocations, small deletions, and junctions passing all filters.

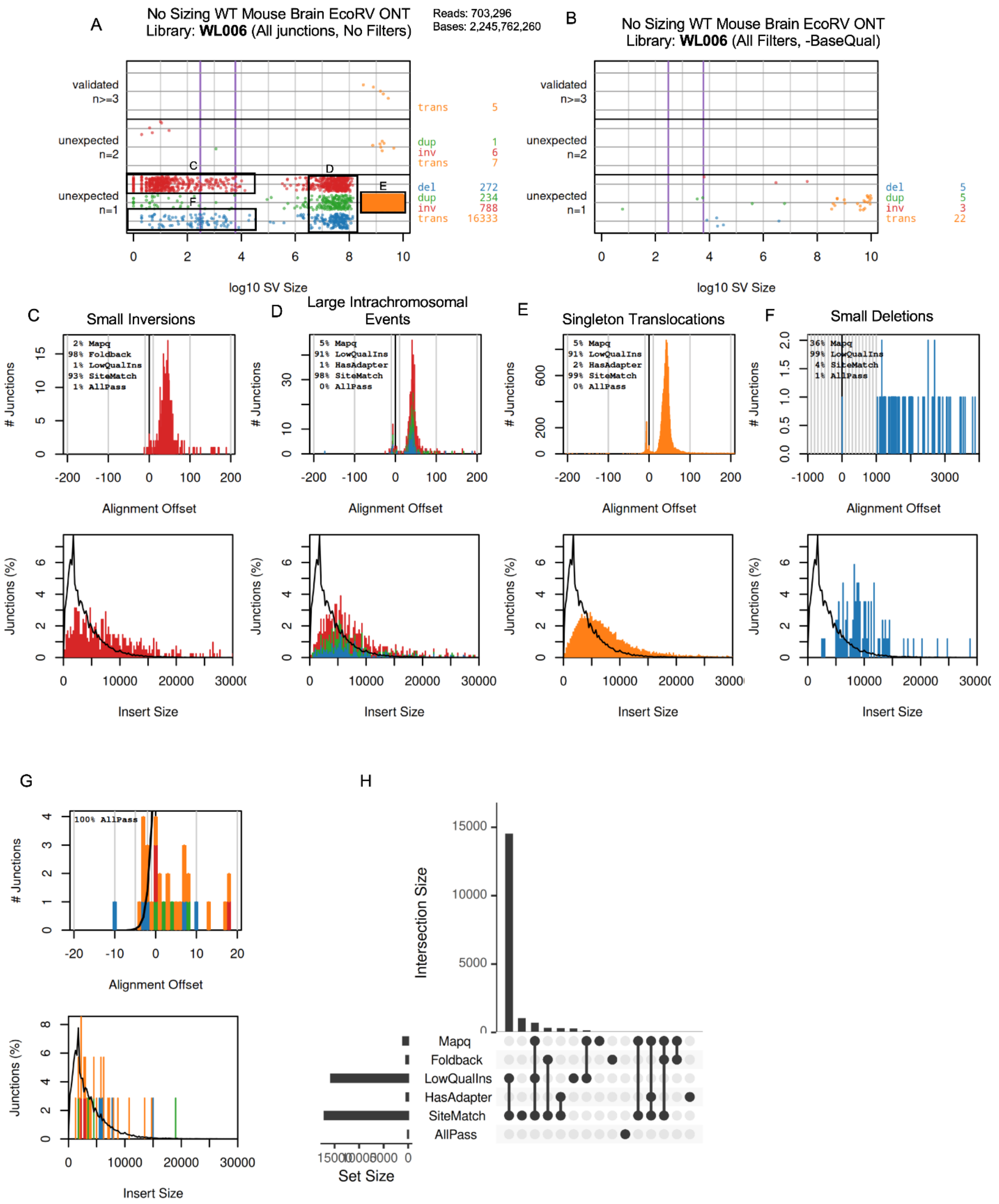


**Figure S4. WL006 mouse brain library complete artifact analysis**

**A.** Junction summary plot like Figure 4A for a non-size-selected mouse brain ONT library without any active filters. Black boxes depict read subsets used in panels C through F.

**B.** Like A, but with all filters active except BaseQual.

**C. to F.** Histograms showing the number of junctions with insertions (positive alignment offset) or microhomology (negative alignment offset) (top) and insert sizes (bottom) for small inversions, large intrachromosomal events, singleton translocations and small deletions. Insets show junction failure percentages by filter type.

**G.** Alignment offset plot (top) and insert size plot (bottom) of remaining junctions when all filters (except BaseQual for nanopore) are applied.

**H.** Upset plot showing the relationship between different filtering flags for all library junctions.

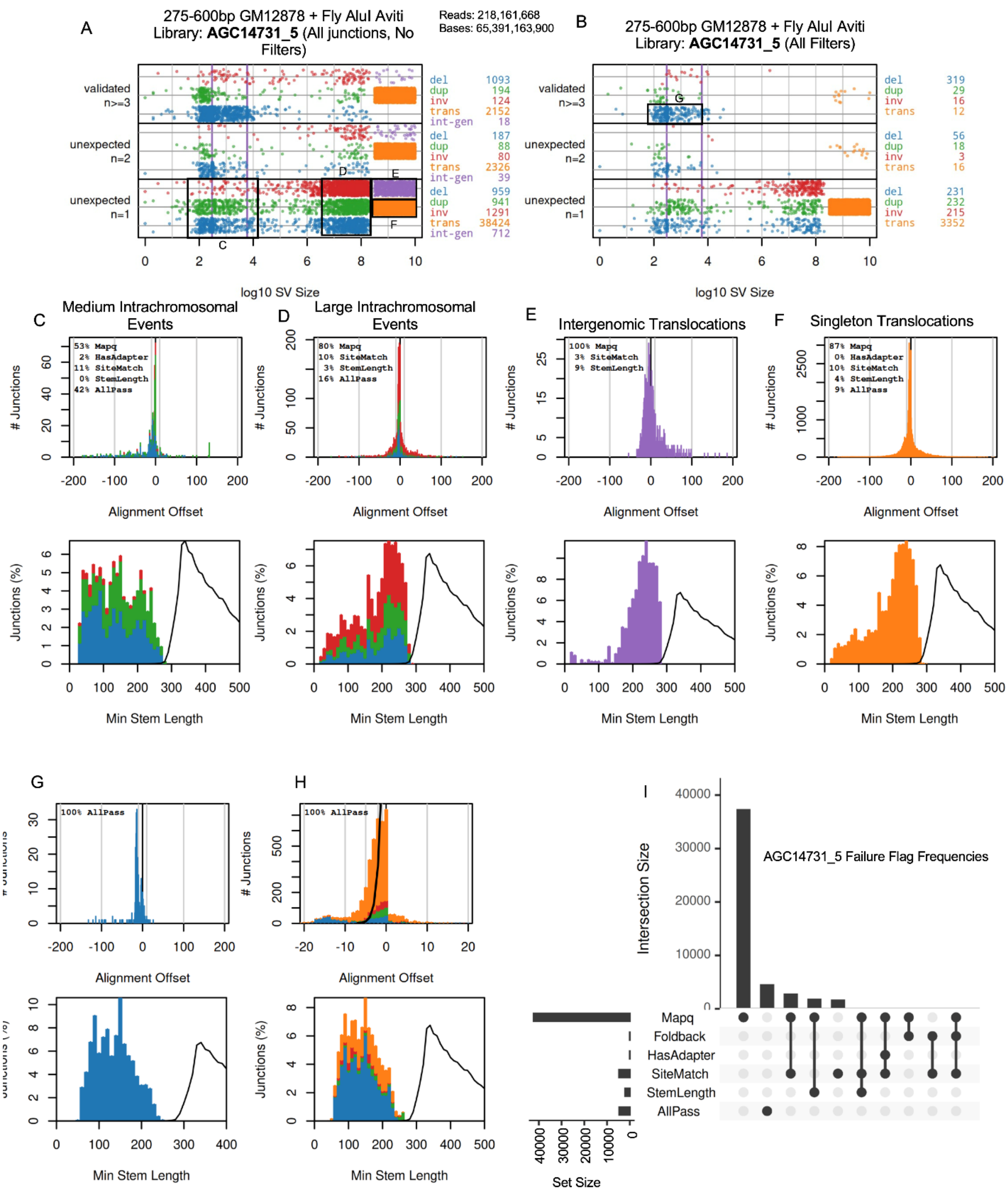


**Figure S5. AGC14371_5 Aviti library complete artifact analysis without MapQ filtering**

**A.** Junction summary plot like Figure 4A for a non-size-selected human and fly Aviti 1x300 library without any active filters. Black boxes depict read subsets used in panels C through E.

**B.** Like A, but with all filters active. The black box depicts the read subset used in panel F.

**C. to F.** Histograms showing the number of junctions with insertions (positive alignment offset) or microhomology (negative alignment offset) (top) and minimum stem lengths (bottom) for medium intrachromosomal events, large intrachromosomal events, intergenomic translocations, and singleton translocations. Insets show junction failure percentages by filter type.

**G.** Histograms like C through E now showing filtered, validated deletions.

**H.** Alignment offset plot (top) and minimum stem length plot (bottom) of remaining junctions when all filters are applied.

**I.** Upset plot showing the relationship between different filtering flags for all library junctions.

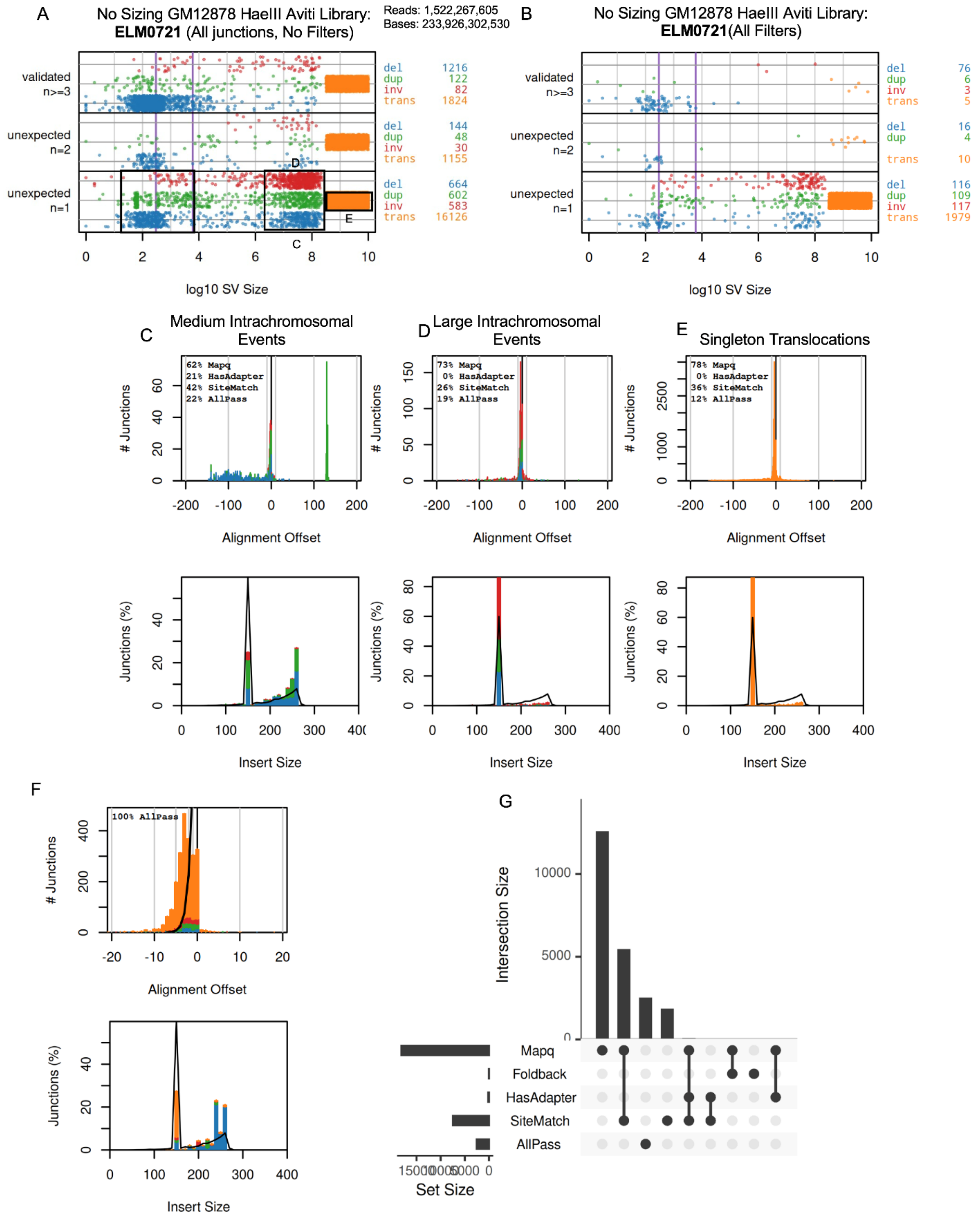


**Figure S6. ELM0721 Aviti library complete artifact analysis**

**A.** Junction summary plot like Figure 4A for a non-size-selected human Aviti 2x150 library without any filters. Black boxes depict read subsets used in panels C through E.

**B.** Like A, but with all filters active.

**C. to E.** Histograms showing the number of junctions with insertions (positive alignment offset) or microhomology (negative alignment offset) (top) and insert sizes (bottom) for medium intrachromosomal events, large intrachromosomal events, and singleton translocations. Insets show junction failure percentages by filter type.

**F.** Alignment offset plot (top) and minimum stem length plot (bottom) of remaining junctions when all filters are applied.

**G.** Upset plot showing the relationship between different filtering flags for all library junctions.

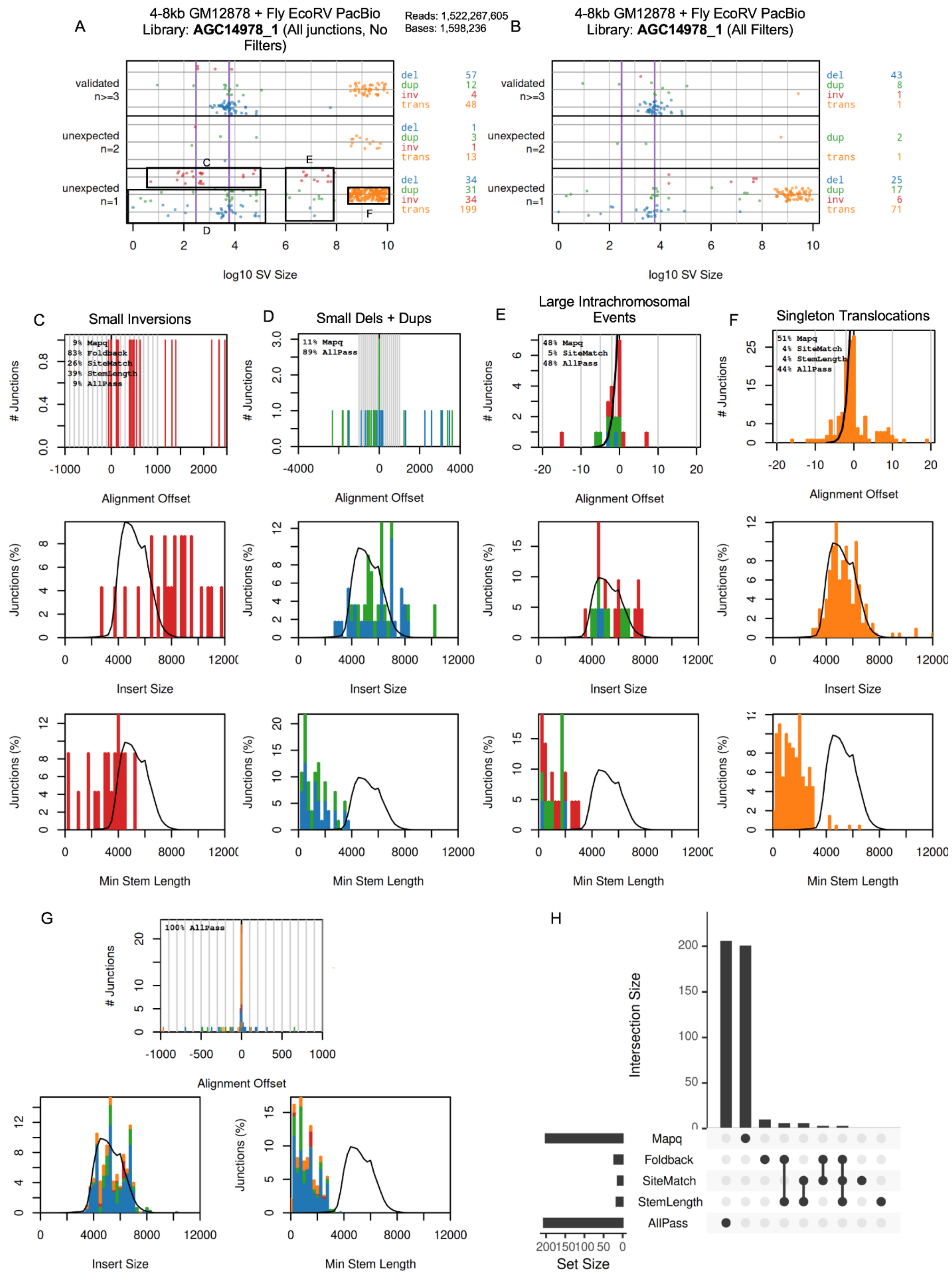
A
4-8kb GM12878 + Fly EcoRV PacBio
Library: AGC14978_1 (All junctions, No Filters)
Reads: 1,522,267,605
Bases: 1,598,236
B
4-8kb GM12878 + Fly EcoRV PacBio
Library: AGC14978_1 (All Filters)
validated n>=3
unexpected n=2
unexpected n=1
log10 SV Size
C
Small Inversions
D
Small Dels + Dups
E
Large Intrachromosomal Events
F
Singleton Translocations
### Junctions
Alignment Offset
Junctions (%)
Insert Size
Min Stem Length
G
100% AllPass
H
Intersection Size
Mapq
Foldback
SiteMatch
StemLength
AllPass
Set Size

**Figure S7. AGC14978_1 PacBio library complete artifact analysis**

**A.** Junction summary plot like Figure 4A for a 4-8kb human + fly PacBio library without any active filters. Black boxes depict read subsets used in panels C through F.

**B.** Like A, but with all filters active.

**C. to F.** Histograms showing the number of junctions with insertions (positive alignment offset) or microhomology (negative alignment offset) (top), insert sizes (middle), and minimum stem lengths (bottom), for small inversions, small deletions and duplications, large intrachromosomal events, and singleton translocations.

**G.** Alignment offset plot, insert size plot, and minimum stem length plot of remaining junctions when all filters are applied.

**H.** Upset plot showing the relationship between different filtering flags for all library junctions.

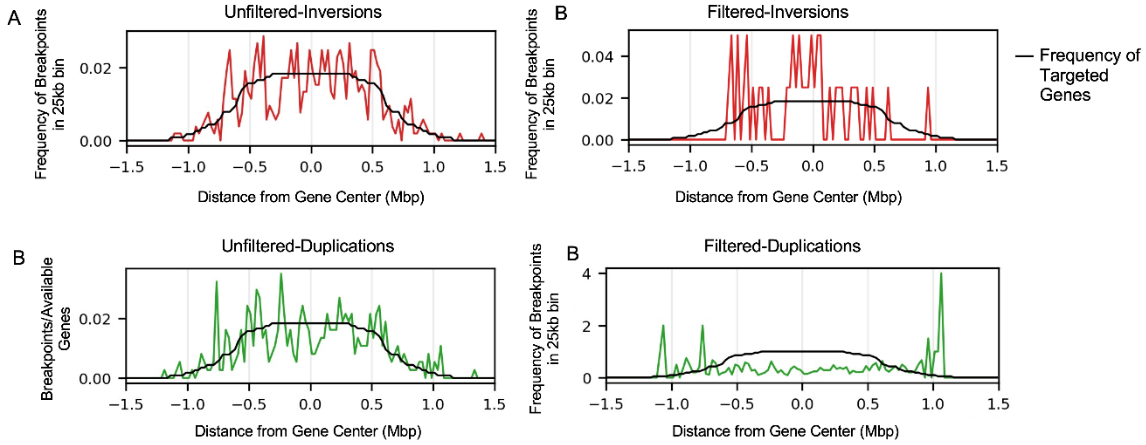


**Figure S8. RPE-1 experiment breakpoint traces for inversions and duplications**

**A. and B.** Traces like Figure 6C,D depicting the RPE-1 inversion breakpoint distribution in 25kb bins without (A) and with (B) error correction filtering. The black line is the frequency distribution of the number of genes targeted by adaptive sampling for the same bins.

**C. and D.** Like A and B, for duplication junctions.

| Mouse Name | Background | Genotype | Tissue | Sequencing Runs |
|---|---|---|---|---|
| B499 | Female B6 | WT | Hippocampus, cortex | WL003_fast, WL003_HAC, WL005, WL006 |
| B502 | Female B6 | WT | Cortex | WL008, WL009, WL010, WL012 |
| Sekiguchi | Male B6/129 | p53-/- | Mixed | WL019, WL021, WL023 |
| JAX | Male B6 | p53-/- | Hippocampus, cortex | WL024, WL025, WL026, WL027, WL029, WL030 |

**Supplementary Data Table 1. Mouse sample information**

**Supplementary Data Table 2. HiFiRe3 library information**

See accompanying Excel file.
